# Synaptic adhesion molecule signaling is activated and organized by tyrosine phosphorylation-induced biomolecular condensate formation

**DOI:** 10.64898/2026.08.25.747070

**Authors:** Shenghan Wu, Nicole A. Morales, Daniel R. Li, Nathan A. McDonald

## Abstract

The precise formation of synapses ensures the proper wiring and function of nervous systems. Specific synapse formation is controlled by synaptic adhesion molecules, which link pre- and post-synaptic cells. Despite this central role, details of how adhesion molecules organize and signal intracellularly to build core synaptic structures are limited. Here, we identify multiple tyrosine phosphorylation sites on the cytoplasmic tail of the *C. elegans* SYG-1 synaptic adhesion molecule that are critical to initiate presynapse formation. We determine that SRC-1 and SRC-2 tyrosine kinases are redundantly responsible for SYG-1 phosphorylation and are consequently critical for presynapse assembly. The phosphorylated population of SYG-1 localizes in clusters within a larger SYG-1 pool and these clusters mark sites of presynaptic active zone assembly. Reconstitution of SYG-1 clusters *in vitro* with SH2-domain adapters and WSP-1 reveals a dynamic biomolecular condensate-forming system. Blocking phosphotyrosine adapters and condensate formation *in vivo* results in the loss of SYG-1 clusters, defective presynapse formation, and compromised neurotransmission. We conclude that phosphorylation of a subpopulation of synaptic adhesion molecules activates and organizes them into condensate-based clusters to initiate presynapse formation.

## Introduction

Nervous system development coordinates the precise wiring of neurons into circuits through synaptic connections^1^. Synapses enable the rapid transfer and transformation of information between neurons. The location and properties of synapses ultimately determine the function of neural circuits that build a nervous system.

Chemical synapses of all types adopt a stereotyped structure, with a presynaptic terminal directly apposed to a postsynaptic specialization, separated by the synaptic cleft. The presynaptic terminal assembles a core active zone structure that organizes calcium channels and release machinery as well as a synaptic vesicle pool^2,3^. The postsynapse forms a postsynaptic density (PSD) that organizes neurotransmitter receptors^4^. Each of these structures have been shown to undergo liquid-liquid phase separation to form biomolecular condensates^5^, concentrated assemblies of protein de-mixed from the cytoplasm^6,7^. Core active zone scaffold proteins SYD-2/Liprin-α, ELKS, RIM, and RIM-BP drive active zone condensate formation^8–11^. Synapsin and synaptophysin further encompass the synaptic vesicle pool into a liquid condensate^12,13^. Postsynaptic scaffolds, including PSD-95 and SynGAP, phase separate to form condensates critical to PSD formation and function^14–16^. Formation of synaptic condensates is critical to build synaptic structures, and the properties of the condensates have an impact on synaptic function^17^.

The core synaptic structures form at specific subcellular sites and between specific cells. Synaptic cell adhesion molecules (SAMs) are hypothesized to act upstream of synapse assembly and direct the process^1,18^. SAMs bind between the synapsing cells and likely signal in both directions to form and shape synapses. A large variety of synaptic cell adhesion molecules capable of selecting proper partner cells and initiating synapse formation have been identified, including Neurexins^19^, Neuroligins^20^, Teneurins^21^, Latrophilins^22^, LAR-type receptor phosphotyrosine phosphatases (LAR-RPTPs)^23^, Ephrin and Eph receptors^24^, a number of Ig-domain adhesion molecules^25,26^, and others. These molecules bind through extracellular interaction domains to confer specificity in synaptic partner choice and control the subcellular location of synapse formation. The great diversity of SAMs creates an incredibly complex interactome^1^. For instance, LAR-RPTPs interact with several different postsynaptic CAMs including TrkC^27^, synaptic adhesion-like molecules (SALMs)^28^, Slit-trks^29^, and Netrin-G ligand 3^30^. Further, different SAMs are specific to certain cell types and synapses^31^. Thus, a complex code of SAMs orchestrates synaptic wiring.

While considerable progress has been made on SAM extracellular interactions driving synaptic specificity, how these molecules subsequently signal intracellularly to direct core synapse structure formation is less clear^1^. A network of intracellular protein-protein interactions has been mapped from certain CAMs, such as presynaptic LAR-PTP^32^. These interactions impact synaptic structure and function, but it is less clear if they drive initial synapse formation. Recent studies also suggest a direct role for certain SAMs in tethering condensate-based synaptic structures. The intracellular domain of presynaptic Teneurin3 is capable of incorporation into RIM and RIM-BP condensates^33^, suggesting a tethering mechanism *in vivo*. Similarly, postsynaptic Latrophilin3 incorporates into PSD-95 condensates^34^. Further, SAMs such as latrophilins are GPCRs that produce cAMP, which may locally signal for synapse formation^35^. It is not yet clear if there are universal SAM signaling mechanisms sufficient for synapse formation; however, synaptic core structures are formed regardless of the exact initiating SAM.

Here, we determine the mechanism initiating cytoplasmic signaling of the *C. elegans* SYG-1 synaptic adhesion molecule. We identify tyrosine phosphorylation within SYG-1’s cytoplasmic tail that induces biomolecular condensate formation through SH2-domain adapter proteins and the Arp2/3 activator WSP-1. Condensate formation organizes the active subpopulation of SYG-1 into clusters to initiate presynapse formation.

## Results

### SYG-1 initiates presynapse formation through phosphorylation of tyrosine residues in its intracellular tail

The *C. elegans* Hermaphrodite Specific Neuron (HSN) forms stereotyped synapses with vulval muscles cells and VC interneurons to control the worm’s egg-laying circuit (Figure 1A)^36–38^. Synapse formation in HSN is controlled by two Ig-domain synaptic adhesion molecules, SYG-1 (Figure 1B) and SYG-2^39,40^. Through extracellular Ig domains, these molecules specifically bind^41^ to specify the HSN synaptic zone and initiate synapse formation. We reasoned that intracellular signaling to initiate the formation of core synaptic structures must originate from the cytoplasmic tail of these adhesion molecules. We first confirmed that the cytoplasmic tail of presynaptic SYG-1 is required for its function by endogenously deleting the domain (Figure 1C). We imaged HSN presynapse formation with endogenous fluorescent markers of the active zone (SYD-2/Liprin-α) and synaptic vesicles (RAB-3). We found a significant loss of synapses in the *syg-1(tailΔ)* mutant, comparable to a loss-of-function *syg-1Δ* allele (Figure 1C-D). The *syg-1(tailΔ)* mutant localized normally to the synaptic zone of HSN, confirming intracellular functions alone were compromised (Figure S1).

**Figure 1:**
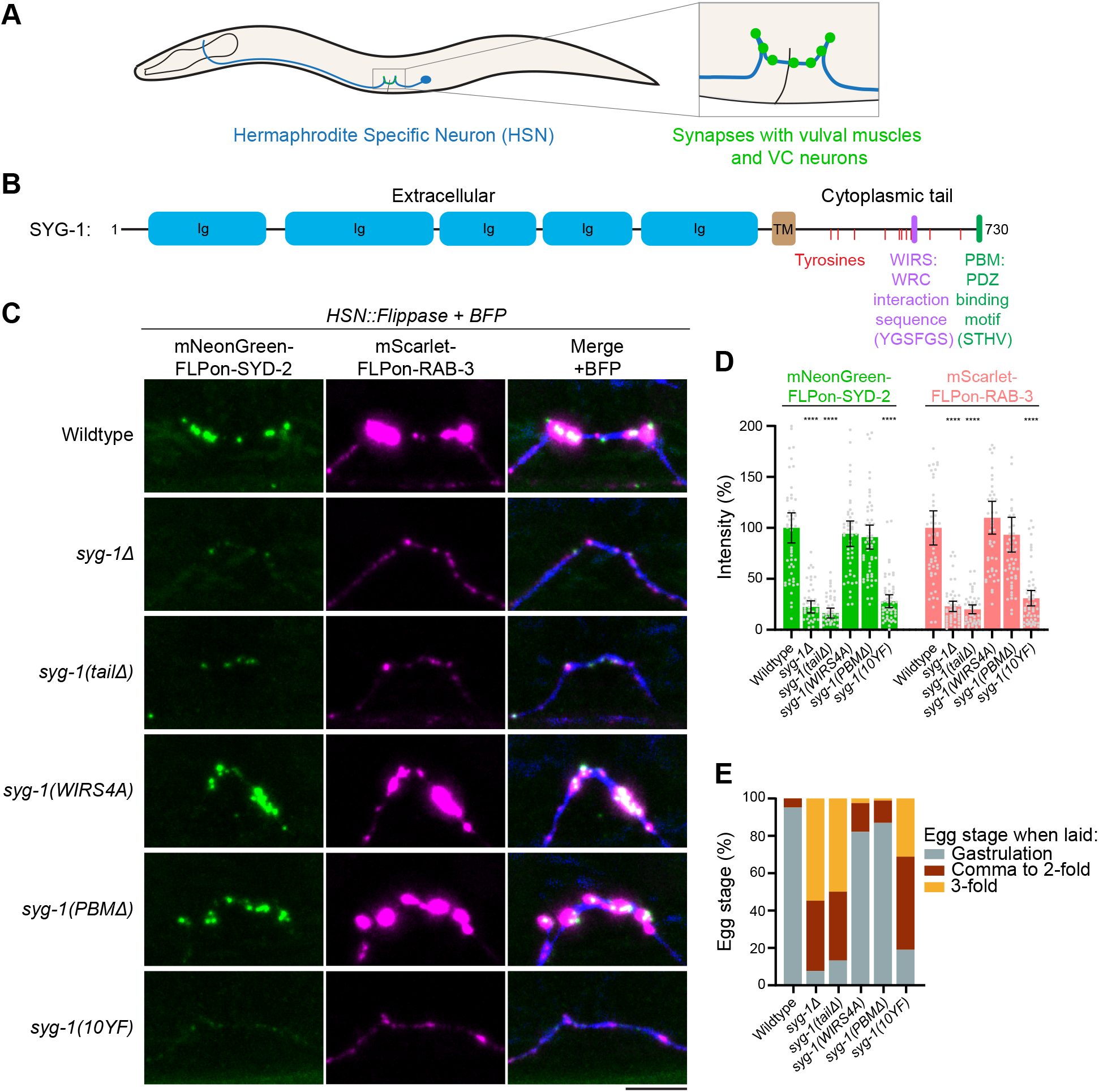
SYG-1 initiates presynapse formation through tyrosine residues in its intracellular tail. A) Schematic of the C. elegans Hermaphrodite Specific Neuron (HSN), which forms synapses with vulval muscles and VC neurons to control egg laying. B) Schematic of the SYG-1 synaptic adhesion molecule with cytoplasmic tail motifs highlighted. C) HSN synapse formation phenotypes in the indicated syg-1 mutants with endogenously-tagged SYD-2 (active zone) and RAB-3 (synaptic vesicles) markers. BFP is expressed cytoplasmically. Scale bar, 5 μm. D) Quantification of HSN synaptic marker fluorescent intensities in (C). ****, p < 0.0001. E) Egg laying assay with the indicated syg-1 mutants. Eggs laid by day 1 adults over 1 hour were classified into gastrulation, comma to 2-fold, and 3-fold stages. Advanced egg development at laying indicates defective HSN circuit function. N > 300 for each genotype.

SYG-1’s cytoplasmic tail is 155 amino acids long and contains two known protein binding motifs: a Wave Regulatory Complex (WRC) interacting receptor sequence (WIRS)^42^ and a predicted PDZ domain binding motif (PBM)^39^ (Figure 1B). We also noticed the tail contains an abundance of tyrosine residues, reminiscent of adhesion molecules and receptors in other systems that are tyrosine phosphorylated for activation^43^ (Figure 1B). To test the importance of these motifs on synapse formation, we endogenously mutated each one to impair its function: a 4 residue alanine substitution in the WIRS site to block WRC interaction^42^, a deletion of the PBM, and a 10YF substitution mutating all tail tyrosines to phenylalanines. Subsequent imaging of HSN synapse formation revealed limited impacts of the WIRS or PBM disruptions^44^; however, a *syg-1(10YF)* mutant showed a strong loss of synapses, comparable to *syg-1(tailΔ)* or *syg-1Δ* alleles (Figure 1C-D). To confirm the impact of the observed synapse formation defects on neurotransmission and HSN circuit function, we performed an egg-laying assay. Compromised HSN synaptic transmission results in stalled egg laying^45^, with eggs advancing far in development before release. Wildtype animals release eggs during the eggs’ gastrulation, while *syg-1Δ*, *syg-1(tailΔ),* and *syg-1(10YF)* mutants showed a significant delay (Figure 1E). The *syg-1(WIRS4A)* and *syg-1(PBMΔ)* alleles displayed significant but minor impacts as well. Together, these data indicate tyrosines in the SYG-1 tail are required for proper HSN synapse formation and function.

To determine if tyrosine residues in the SYG-1 tail are in fact phosphorylated, we isolated SYG-1 endogenously fused with a SNAP tag for analysis with mass spectrometry. O^6^-benzylguanine beads were used to covalently bind the tagged SYG-1, subsequently enabling the use of stringent and denaturing conditions to extract it from stable and detergent resistant synaptic structures in its phosphorylated form (Figure 2A-B). Phosphopeptide enrichment was further necessary, as phosphorylation events were scarce and not detected in a direct SYG-1 pulldown sample. After enrichment, mass spectrometry analysis revealed 7 phosphotyrosine sites in the SYG-1 tail (Y603, Y608, Y623, Y649, Y660, Y686, and Y711) (Figure 2C, Figure S2, and Table S1). We constructed a 7YF mutant to determine if these sites were sufficient to reproduce the synaptic loss phenotype seen in the 10YF mutant. Indeed, a *syg-1(7YF)* mutant displayed a similar defect in synapse formation seen in *syg-1(10YF)* (Figure 2D and 2E) and consequent HSN circuit function (Figure 2F), indicating that these 7 phosphotyrosines are required for SYG-1 tail signaling. Together, these data confirm that multiple tyrosine phosphorylation sites in SYG-1’s tail are critical for SYG-1’s synapse formation function.

**Figure 2:**
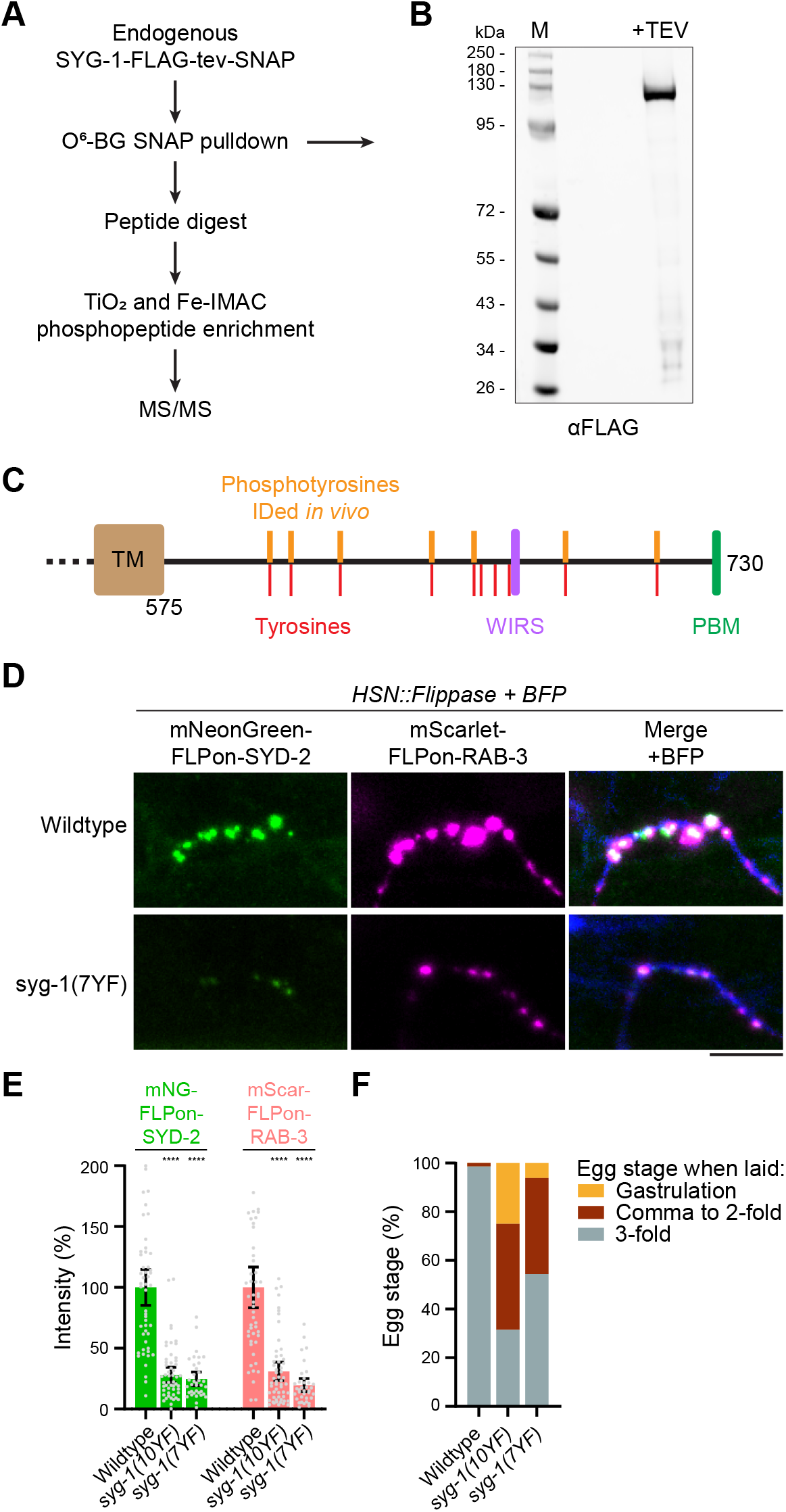
The SYG-1 intracellular tail is tyrosine phosphorylated to initiate synapse formation. A) Workflow of a SYG-1 SNAP-tag pulldown and phosphopeptide enrichment. O6-BG: O6-benzylguanine; TiO2: titanium dioxide; Fe-IMAC: Iron immobilized metal ion affinity chromatography. B) Western blot of SYG-1-FLAG-tev-SNAP pulldown. TEV protease digestion was performed to release the covalent SNAP tag link to O6-BG beads. C) Schematic of SYG-1 intracellular tail with phosphotyrosines identified by mass spectrometry (Y603, Y608, Y623, Y649, Y660, Y686, and Y711). See Figure S2 and Table S1 for spectra. D) HSN synapse formation phenotypes in the indicated syg-1 mutants with endogenously-tagged SYD-2 (active zone) and RAB-3 (synaptic vesicles) markers. BFP is expressed cytoplasmically. Scale bar, 5 μm. E) Quantification of HSN synaptic marker fluorescent intensities in (D). ****, p < 0.0001. F) Egg laying assay with the indicated syg-1 mutants. Eggs laid by day 1 adults over 1 hour were classified into gastrulation, comma to 2-fold, and 3-fold stages. Advanced egg development at laying indicates defective HSN circuit function. N > 300 for each genotype.

### Proximity labelling identifies SYG-1 intracellular tail kinases and interactors

To investigate what kinases are responsible for SYG-1 phosphorylation and identify possible binding partners and effectors of this signal, we performed proximity labeling on SYG-1’s cytoplasmic tail^46^. We endogenously tagged the C-terminal end of SYG-1’s cytoplasmic tail with TurboID (Figure 3A), treated animals with biotin, and pulled down the resulting biotinylated proteins with streptavidin beads. Subsequent mass spectrometry analysis identified 1311 proteins significantly enriched versus a control streptavidin pulldown (Figure 3B and Table S2). Gene ontology analysis revealed an enrichment of proteins involved in cytoskeletal organization, axon development, cell-cell junction formation, and other processes consistent with previously described SYG-1 functions in synapse formation and axon branching^39,42^ (Figure 3C). The identified proteins were also enriched for intriguing protein domains related to these functions, including tyrosine kinase, SH2 and phosphotyrosine-binding, SH3, and PDZ domains (Figure 3D).

**Figure 3:**
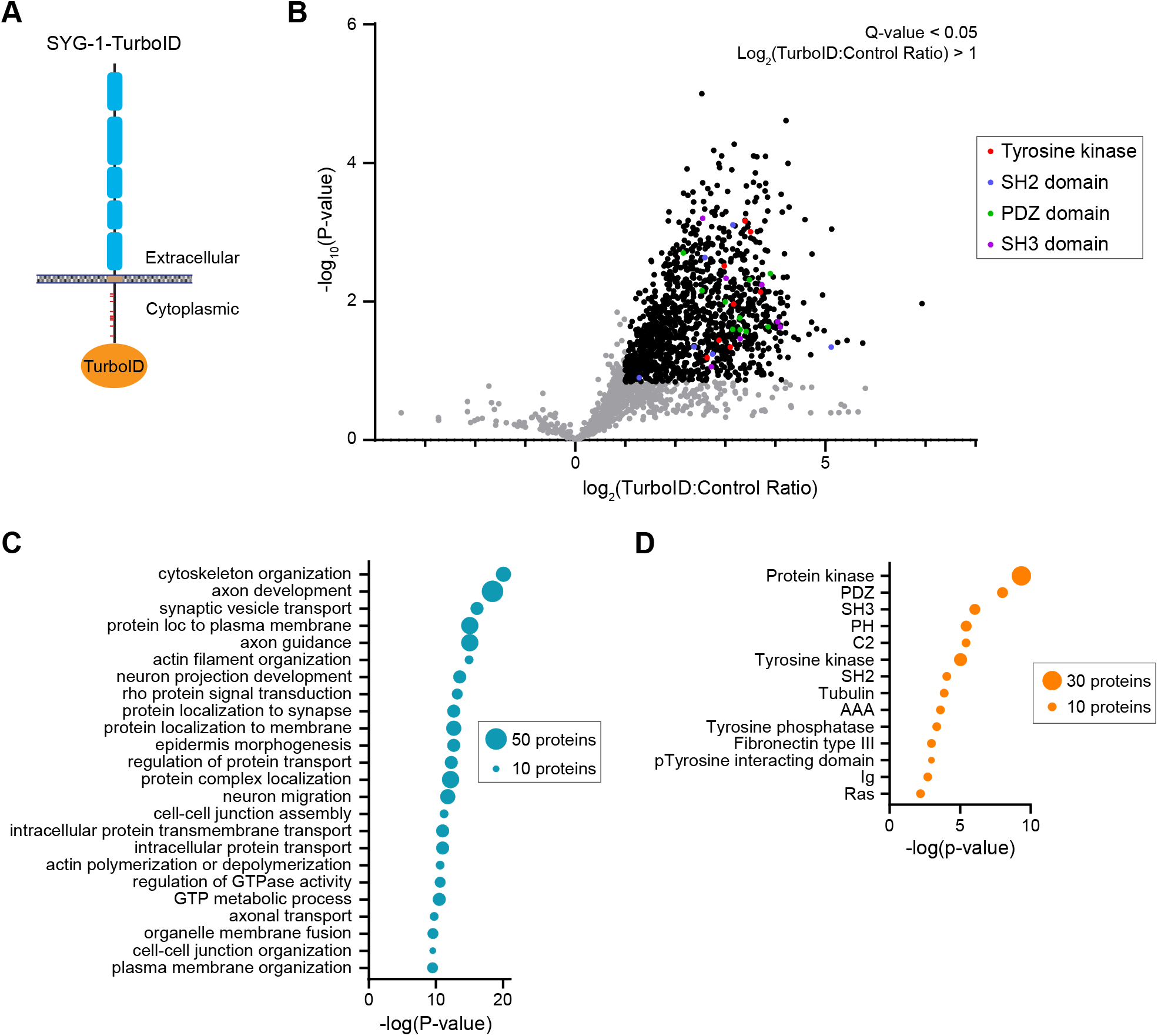
TurboID-mediated proximity labelling identifies SYG-1 intracellular tail interactors. A) Schematic of endogenously-tagged SYG-1-TurboID. B) Proteins identified by mass spectrometry in SYG-1-TurboID samples versus an untagged control. Bold dots are proteins considered significant with log2(TurboID:Control ratio) > 1 and Q-value < 0.05. Proteins containing functional domains of interest are highlighted. C, D) Gene Ontology (GO) enrichment analysis of significantly represented biological processes (C) and protein domains (D) in the identified interactors. Dot size represents the number of genes matching the category. See Table S2 for full data.

### SRC-1 and SRC-2 redundantly phosphorylate the SYG-1 intracellular tail and are critical for synapse formation

We next sought to identify the tyrosine kinases responsible for SYG-1 tail phosphorylation from TurboID interactors (Table S2). We tested null mutants of the identified tyrosine kinases for impacts on HSN synapse formation. We found a *src-2Δ* mutant showed a modest synapse formation phenotype (Figure 4A-B). A *src-1Δ src-2Δ* double mutant showed a strong loss of HSN synapses (Figure 4A-B), implicating these two kinases in synapse formation perhaps through SYG-1 phosphorylation. To determine if these kinases directly phosphorylated SYG-1, we performed *in vitro* kinase assays with purified recombinant SYG-1 cytoplasmic tail and SRC-1 and SRC-2 kinases (Figure 4C). These assays revealed robust phosphorylation of the SYG-1 tail by both SRC-1 and SRC-2. The kinases also autophosphorylated (Figure 4C), consistent with Src family kinase regulation^47,48^. Mass spectrometry analysis of the *in vitro* phosphorylated SYG-1 tail revealed 8 of 10 tyrosines were phosphorylated by both SRC-1 and SRC-2 (Figure S3A and Table S3), overlapping with the sites found *in vivo*. Kinase assays with a SYG-1(10YF) mutant abolished phosphorylation *in vitro* (Figure S3). Together, these data indicate that SRC-1 and SRC-2 kinases redundantly phosphorylate the SYG-1 cytoplasmic tail and are consequently critical for HSN synapse formation.

**Figure 4:**
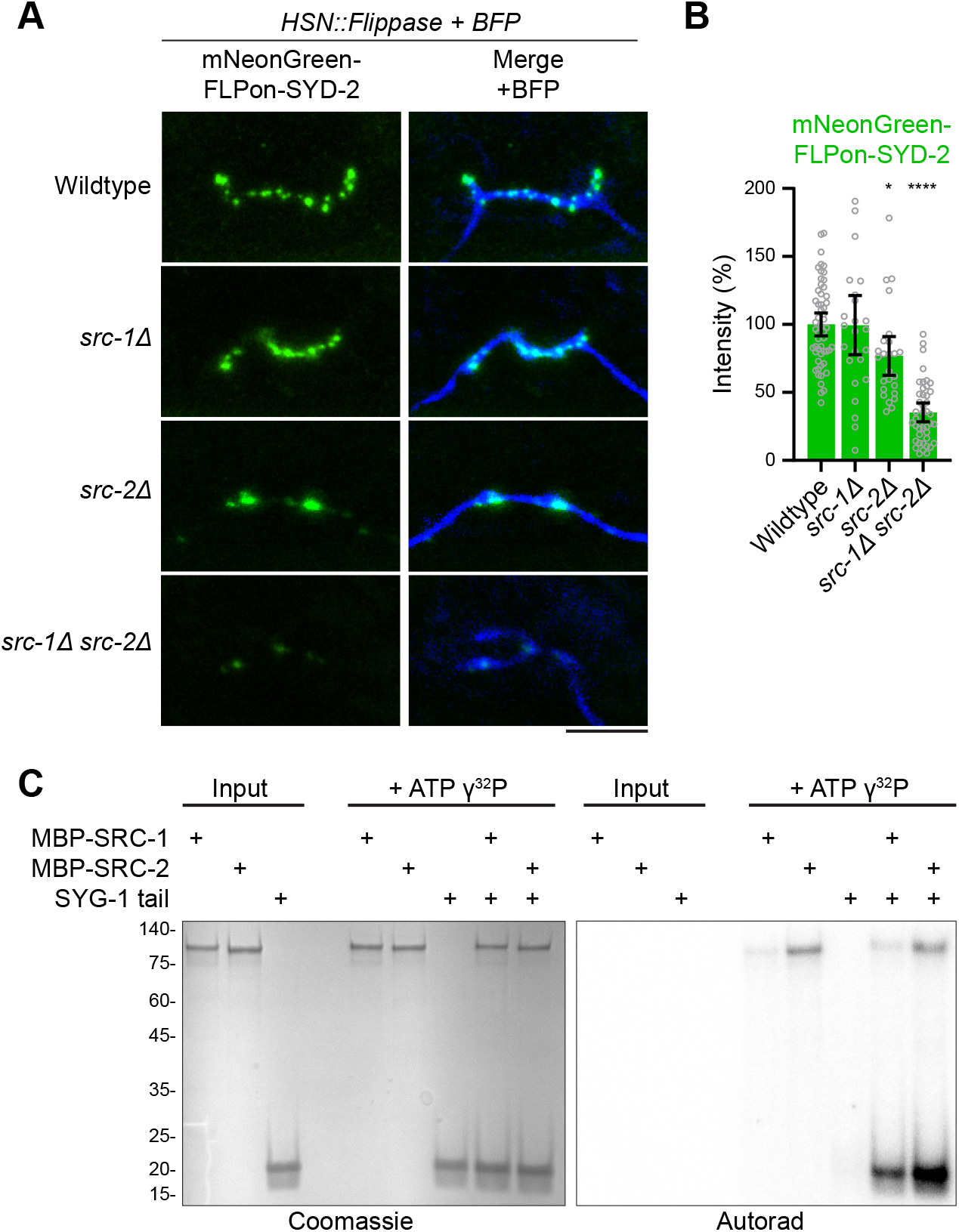
SRC-1 and SRC-2 tyrosine kinases redundantly phosphorylate the SYG-1 intracellular tail and are critical for synapse formation. A) HSN synapse formation phenotypes in the indicated mutants with an endogenously-tagged SYD-2 active zone marker. BFP is expressed cytoplasmically. Scale bar, 5 μm. B) Quantification of HSN synaptic marker fluorescent intensities in (A). *, p < 0.05; ****, p < 0.0001. C) In vitro kinase assay between purified recombinant SRC-1, SRC-2, and the SYG-1 tail.

### A subpopulation of phosphorylated SYG-1 forms clusters coincident with active zone formation

With the identification of critical phosphosites in SYG-1 and the responsible kinases, we next sought to visualize where the phosphorylated SYG-1 species are found in HSN. Methods traditionally used to probe localization of phosphorylated proteins such as phospho-specific antibodies are challenging to apply in *C. elegans.* We therefore sought a genetically encodable sensor for SYG-1 phosphorylation. Proximity labeling experiments identified multiple potential interacting proteins containing SH2 domains, known to directly bind phosphotyrosine sites^49,50^. We therefore engineered a SYG-1 phosphotyrosine sensor by fusing isolated SH2 domains from these interactors with a fluorophore for visualization. We found the SH2 domain from NCK-1, a known interactor of SYG-1’s human orthologs Neph1 and Nephrin^51,52^, effectively marked phosphorylated SYG-1 in HSN (Figure 5A). This sensor was localized to small puncta within the HSN synaptic zone and localization was lost in the *syg-1(10YF)* mutant, confirming specificity to SYG-1 phosphotyrosine sites (Figure 5A-B). Importantly, since the SH2 domain is likely to directly bind phosphorylated SYG-1 and mask functional phosphosites, we investigated if sensor expression altered the formation of synapses, and found no impact (Figure S4).

**Figure 5:**
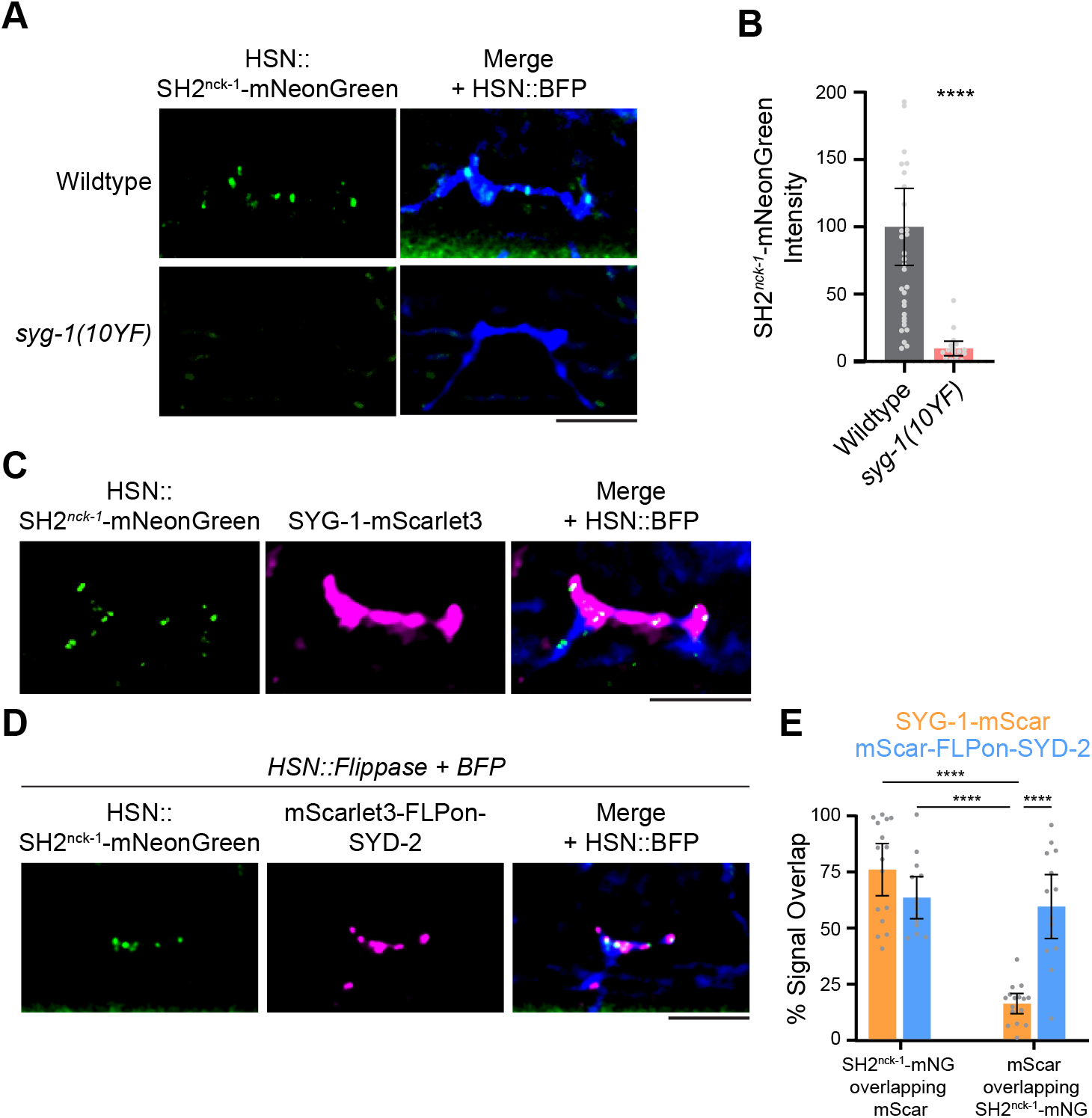
A subpopulation of phosphorylated SYG-1 forms clusters coincident with sites of active zone formation. A) Confocal microscopy of an SH2nck-1-mNeonGreen sensor in wildtype and a syg-1(10YF) mutant. BFP is expressed cytoplasmically. The lack of sensor localization in the non-phosphorylatable syg-1(10YF) mutant indicates it is specific for SYG-1 tyrosine phosphorylation. B) Quantification of SH2nck-1-mNeongreen intensity in (B). ****, p < 0.0001. C) Confocal microscopy of an SH2nck-1-mNeonGreen SYG-1 phosphorylation sensor and endogenous SYG-1-mScarlet3 in HSN. BFP is expressed cytoplasmically. A small subpopulation of clustered, phosphorylated SYG-1 is present within the larger SYG-1 localization zone. D) Confocal microscopy of an SH2nck-1-mNeonGreen sensor and an endogenous mScar-let3-FLPon-SYD-2 active zone marker in HSN. BFP is expressed cytoplasmically. E) Quantification of overlap between SH2nck-1-mNeonGreen, SYG-1-mScarlet3, and mScarlet3-FLPon-SYD-2 in (A) and (D). Most active zones marked by SYD-2 overlap with phosphorylated SYG-1 clusters, while these clusters represent only a small subpopulation of the total SYG-1. ****, p < 0.0001. Scale bars, 5 μm.

We next visualized this SYG-1 phosphorylation sensor together with labelled SYG-1-mScarlet3. SYG-1 is localized to the synaptic zone on the “plateau” of the HSN axon through binding its partner SYG-2^40^ (Figure 1A). A large amount of SYG-1 is present across the membrane in this region (Figure 5C and Figure S1). The SH2^nck-1^-mNeonGreen sensor revealed that phosphorylated SYG-1 is a small subpopulation that localizes to discrete clusters within the large SYG-1 pool (Figure 5C). This stoichiometry explains our challenge of recovering phosphopeptides in a SYG-1 pulldown (Figure 2A-C). We next compared the localization of the phosphorylated SYG-1 clusters with sites of active zone formation marked by SYD-2 and found a strong colocalization (Figure 5D). Quantification of the overlap revealed a majority of SYD-2 active zone sites overlap with a cluster of phosphorylated SYG-1 (Figure 5E). Therefore, these data suggest a small subpopulation of SYG-1 is phosphorylated and forms clusters that mark the sites of active zone formation.

### Phosphorylated SYG-1 tail and partners form biomolecular condensates critical for synapse formation

We next sought to determine how phosphorylated SYG-1 clusters are formed and test if these clusters are critical for synapse formation. Given the effective labeling of phosphorylated SYG-1 by the SH2^nck-1^-mNeonGreen sensor (Figure 5A-D) and SH2 proteins’ crucial roles in other phosphotyrosine signaling pathways^53,54^, we investigated SH2-domain containing proteins as possible adapters and effectors of SYG-1 phosphorylation. From proximity labeling data, we identified multiple potential SH2 domain interactors (Figure 3C and Table S2). We performed *in vitro* binding assays between phosphorylated SYG-1 tail and SH2 domains from identified interactors to reveal three that directly bound: NCK-1, SEM-5/Grb2, and RIN-1 (Figure 6A). These interactions were specific to SYG-1 tail species phosphorylated by SRC-1 and SRC-2, with no interaction observed in the unphosphorylated state.

**Figure 6:**
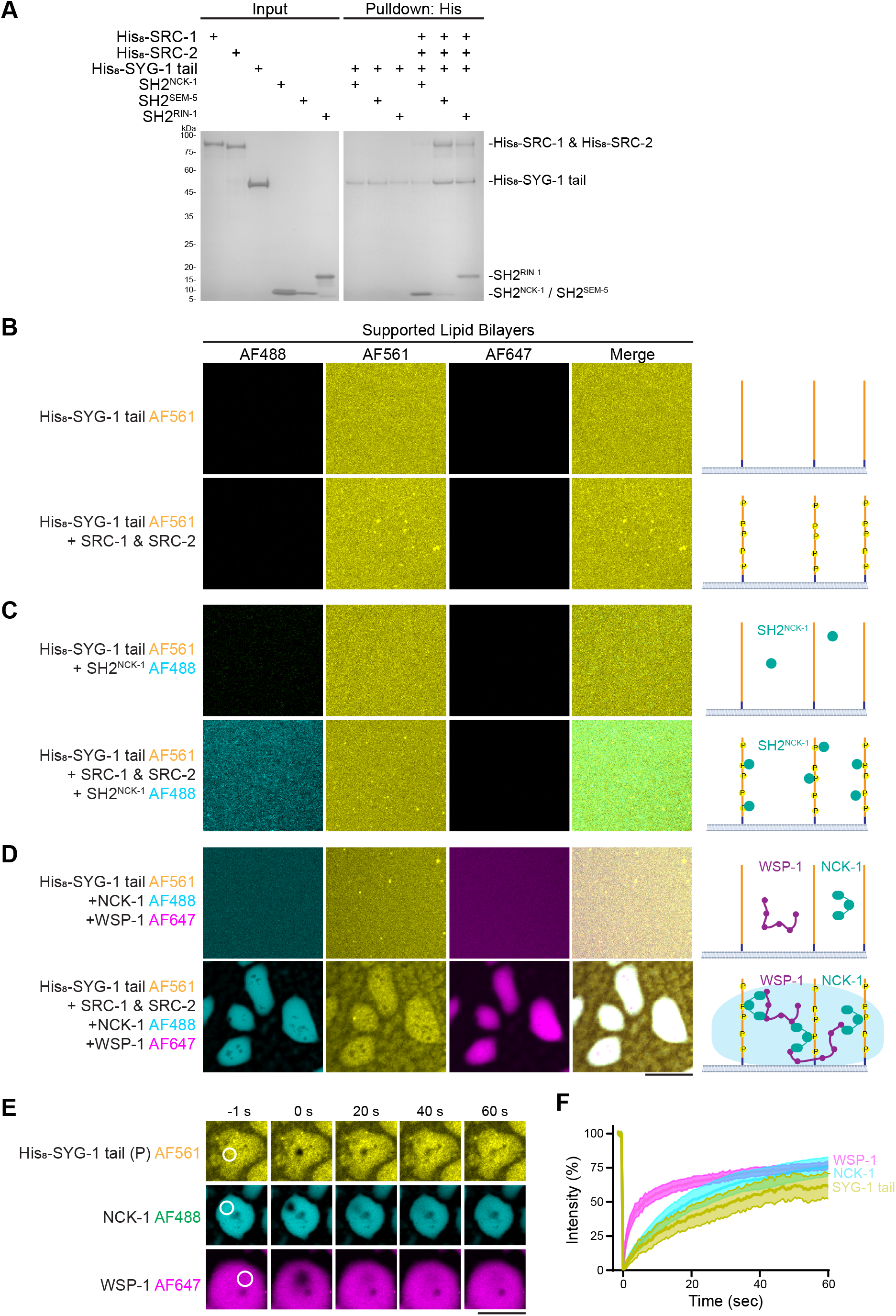
The phosphorylated SYG-1 tail interacts with SH2-domain adapters to form biomolecular condensates. A) In vitro binding assay between the SYG-1 cytoplasmic tail and SH2 adapters identified by proximity labeling. SYG-1 tail was phosphorylated by SRC-1 and SRC-2 overnight before SH2 domains were added. B-D) SYG-1 cytoplasmic tail-bound supported lipid bilayers. B) Addition of SRC-1 and SRC-2 kinases to phosphorylate the SYG-1 tail. C) Addition of the SH2 domain from NCK-1 along with SRC-1 and SRC-2 to SYG-1 tail-bound SLBs reveals phosphorylation. D) Addition of NCK-1 and WSP-1 to the SYG-1 tail-bound SLBs. These partners induce the formation of biomolecular condensates and clustering when SYG-1 is phosphorylated by SRC-1 and SRC-2. E) Fluorescence recovery after photobleaching (FRAP) of each component from (D) reveals liquid dynamics within condensates. The indicated circular white region was bleached and recovery monitored. F) Quantification of FRAP in (E). Scale bars, 10 μm.

In other systems, SH2 adapter proteins, including Nck-1^55,56^ and Grb-2^57–59^, can link with actin network proteins like Wasp, an Arp2/3 activator, or Sos1, a Ras/Rac guanine nucleotide exchange factor (GEF), to form biomolecular condensates^60^. Multivalent interactions that support condensate formation arise from multiple phosphotyrosine binding sites and multiple protein binding domains (often SH3 domains and proline rich motifs) in adapter and effector proteins. Collectively, these interactions induce phase separation and biomolecular condensate formation. In fact, we find *C. elegans* WSP-1 and SOS-1 present in SYG-1 proximity labeling data (Table S2). We hypothesized that a similar mechanism could be at play in SYG-1 synapse formation, which could explain the clustering of phosphorylated species seen *in vivo* (Figure 5A-D). To test this possibility, we reconstituted the SYG-1 tail signaling *in vitro* with recombinant proteins on supported lipid bilayers (SLBs). We assembled purified His_8_-tagged SYG-1 tail on SLBs containing 2% DGS-NTA lipids to reconstitute SYG-1’s normal orientation (Figure 6B and Figure S5A). We then added SRC-1 and SRC-2 kinases and ATP to phosphorylate the SYG-1 tail (Figure 6B). Phosphorylation was confirmed with binding of the SH2 domain from NCK-1 (Figure 6C). Next, we added full-length versions of possible SH2 adapters NCK-1 and SEM-5, as well as WSP-1. Addition of these components to unphosphorylated SYG-1 tail on SLBs resulted in no change (Figure 6D and Figure S5C). However, the combination of phosphorylated SYG-1 tail, either NCK-1 or SEM-5, and WSP-1 resulted in robust formation of biomolecular condensates upon the SLBs (Figure 6D and Figure S5C). Each component in the condensates was found to be highly dynamic when tested with a fluorescence recovery after photobleaching (FRAP) assay (Figure 6E-F and Figure S5D). We were unable to produce full length RIN-1, and addition of its SH2 domain alone was not sufficient for condensate formation (Figure S5E). Confirming the requirement for multivalent interactions, individual additions of NCK-1, SEM-5, or WSP-1 were unable to form condensates (Figure S5F). These data indicate the phosphorylated SYG-1 tail and binding partners are capable of clustering through the formation of biomolecular condensates driven by multivalent interactions.

To determine if this mechanism is important *in vivo*, we deleted the identified SYG-1 adapters to block the formation of condensates. *Nck-1* and *rin-1* were deleted genetically and SEM-5 protein was degraded specifically in HSN with a ZF-degradation system^61^, as it is essential. We first imaged the SH2^nck-1^ SYG-1 phosphorylation sensor in a *nck-1Δ sem-5-ZF* double mutant and a *nck-1Δ sem-5-ZF rin-1Δ* triple mutant and found phosphorylated SYG-1 clusters were abolished (Figure 7A-B). It is likely phosphorylated species still exist which are below the fluorescence detection limit of the sensor. This result confirms SH2 adapter proteins are necessary for SYG-1 condensate cluster formation *in vivo*.

**Figure 7:**
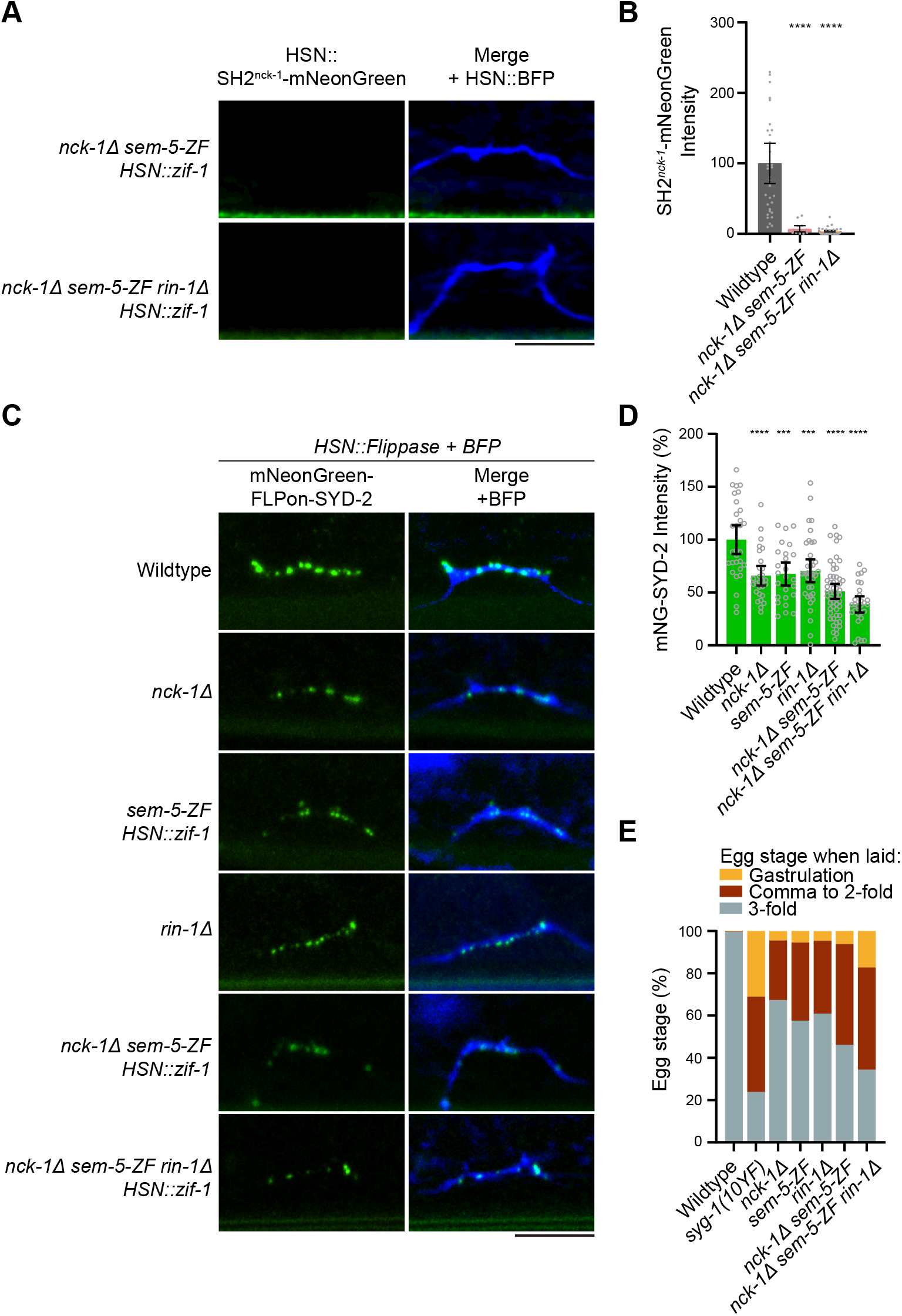
SYG-1 condensate-inducing SH2 domain effectors are critical for HSN synapse formation and function. A) Confocal microscopy of the SH2nck-1-mNeonGreen SYG-1 phosphotyrosine sensor in the indicated mutants. BFP is expressed cytoplasmically. Scale bars, 5 μm. B) Quantification of SH2nck-1-mNeonGreen HSN intensity in (A). ****, p < 0.0001. C) HSN synapse formation phenotypes in the indicated mutants with an endogenously-tagged SYD-2 active zone marker. BFP is expressed cytoplasmically. D) Quantification of HSN synaptic marker fluorescent intensities in (C). ****, p < 0.0001. E) Egg laying assay with the indicated mutants. Eggs laid by day 1 adults over 1 hour were classified into gastrulation, comma to 2-fold, and 3-fold stages. Advanced egg development at laying indicates defective HSN circuit function. N > 300 for each genotype. Scale bars, 5 μm.

Finally, we tested if phosphorylated SYG-1 condensates are important for SYG-1’s function in synapse formation. We imaged HSN synapse formation with a SYD-2 active zone marker in the SH2 domain adapter mutants. *Nck-1Δ, sem-5-ZF*, and *rin-1Δ* alleles were each found to moderately impact HSN synapse formation (Figure 7C-D). Combination mutants further increased the impact to result in a strong loss of synapse formation, similar to our original *syg-1(10YF)* mutation (Figure 7C-D). An egg laying assay further confirmed the synapse formation defects impacted HSN circuit function (Figure 7E). Together, these data indicate phosphorylated SYG-1 condensates formed through SH2-domain adapters are critical for HSN synapse formation and function.

## Discussion

In this study, we have identified cytoplasmic tyrosine phosphorylation of the SYG-1 synaptic adhesion molecule to be critical for its synapse formation function. We determined the SRC-1 and SRC-2 tyrosine kinases are responsible for the phosphorylation and find phosphorylated SYG-1 organizes into clusters *in vivo*. A condensate-based mechanism drives this clustering, induced by binding of SH2-domain adapters and WSP-1. These condensates are critical *in vivo* for synapse assembly and circuit function.

### Condensate-based clustering and activation of adhesion molecules

SYG-1 condensate formation through multivalent binding partners is orthologous to previously characterized Nephrin-based condensates^56^ and immunological synapse-based condensates^57,62^. Condensates formed between phosphorylated Nephrin, SH2 adapter Nck-1, and Wasp help recruit the adhesion molecule in the membrane to clusters that subsequently promote actin polymerization and downstream signaling^63,64^. Three tyrosine residues in Nephrin’s cytoplasmic tail are critical for the signaling^52^. SYG-1 is distantly orthologous to Nephrin, but intracellular tail sequences have diverged between the two, without clear conservation of tyrosine residue sites. However, the mechanism and binding partners appear to be conserved, highlighting the importance of these condensates in each molecule’s function. Similarly, T cell receptor phosphorylation triggers condensate formation through Grb-2 (ortholog of *C. elegans* SEM-5) and Sos1^57^. Thus, condensate-based clustering mechanism appear to be widespread in multiple cellular contexts where receptors and adhesion molecules operate.

Many functional consequences of these condensates have been proposed. Foremost, clustering through condensate formation may amplify active populations of adhesion molecules^55^. Concentrated condensates of Nephrin increase the dwell time of Wasp and Arp2/3 effectors, increasing F-actin assembly output. Further, kinetic proofreading mechanisms are used by the T cell receptor to differentiate transient and long-dwelling interactions, mediated by condensates^59,65^. Here, our evidence suggests that a limited subpopulation of SYG-1 is activated by tyrosine phosphorylation and organized into condensate-based clusters. A large population remains unphosphorylated but localized in the synaptic zone of the axon through binding to its extracellular partner SYG-2. Condensate formation may therefore consolidate the active SYG-1 population into signaling clusters which effectively seed the formation of the presynapse. Why a large population of SYG-1 remains unactivated by phosphorylation is unclear. Through strong binding to its partner SYG-2^41^, it may be important for mechanical stability of the synaptic region of the HSN axon.

### Coordination of multiple synaptic condensates

The synapse has emerged as a hotspot for condensates, including active zone^8,11^, synaptic vesicle pool^12^, and postsynaptic density^15,56^ condensates. With our discovery of synaptic adhesion molecule condensates, it will be critical to determine if and how each interact and how interactions contribute to function. Experiments between RIM and RIM-BP active zone condensates and synaptic vesicle condensates suggest an immiscibility^66^, with synaptic vesicle handover occurring at the boundaries. It is feasible that adhesion molecule condensates could similarly be unable to mix and simply border other condensates. Alternatively, as suggested with Teneurin and Latrophilin^33,34^, these could mix and perhaps seed the formation of core synapse condensates. Direct observation of synaptic condensate formation might resolve these questions, though observation and manipulation of native *in vivo* condensates remains challenging due to their nanometer scale.

### Actin network nucleation in synapse formation

SYG-1’s condensate network identified here contains NCK-1, SEM-5/Grb2, and the Arp2/3 activator WSP-1. We further identified RIN-1, a Ras GEF, as a phosphorylated tail interactor, though we were unable to purify the large protein to test for any condensate forming activity. Previously, the WRC complex was also shown to bind to SYG-1’s tail^42^. These interactors implicate actin network formation as the next step in SYG-1-mediated presynapse formation. Adhesion molecules link to F-actin networks in a variety of contexts^67,68^, suggesting this could be a common pathway.

In fact, multiple actin networks are present at synapses^69,70^. These include branched actin networks at active zones^71^ and post-synaptic densities^72^. A cage of actin further surrounds the synaptic vesicle pool condensates^73^. The developmental timing of these F-actin networks’ formation is not clear. It is possible that F-actin network formation is an initial step in synapse formation downstream of adhesion molecules^74^. Indeed, large scale depolymerization of F-actin networks has a strong negative impact on synapse formation^75^. This raises the question of what connects a SAM-induced actin network to synapse formation? Previous studies in *C. elegans* identified Neurabin NAB-1, an actin-binding protein, to connect F-actin to active zone component SYD-1 and synapse formation^75^. Removal of NAB-1 impacts synapse formation in HSN, but not to the same level of SYG-1, suggesting the involvement of more components. Additional investigation is required to close the link between adhesion molecules, actin networks, and synapse formation.

## Methods

### *C. elegans* methods

*C. elegans* strains (Table S4) were raised on OP50 *Escherichia coli*-seeded nematode-growth medium plates. N2 Bristol was used as the wildtype reference strain. Transgenic strains were created by gonadal microinjection at 50 ng/μL (*Pegl-6*) with 50 ng/μL *Podr-1::RFP or Punc-122::RFP* co-injection markers. Arrays were integrated into the genome with trimethylpsoralen/UV mutagenesis. Egg-laying assays were carried out with day 1 adult hermaphrodites, which were transferred to fresh plates to lay eggs for 1 hour. The developmental stage of the laid eggs was immediately classified into gastrulation, comma to 2-fold, and 3-fold stages.

### Constructs and cloning

All constructs (Table S5) were created with an isothermal assembly method using overlapping oligonucleotides. SYG-1 tail 10YF and 7YF mutant sequences were synthesized (TwistBio) and cloned into a pSK vector. A *C. elegans* codon-optimized 3xFLAG-2xTEV-SNAP and TurboID cassette were synthesized (TwistBio) and cloned into a pSK vector. Three identical SH2 domain sequences from NCK-1 were distinctly codon optimized to enable synthesis with an mNeonGreen sequence and assembled into a pSM vector containing the 3,527-nucleotide promoter of *egl-6.* SRC-1, SRC-2, SYG-1 tail, NCK-1, SEM-5, WSP-1, and RIN-1 sequences were amplified from cDNA and assembled into pET15, pMal-c2, pHis8-tev, and pHis-MBP-tev vectors for bacterial expression. The *zif-1* gene was amplified from cDNA and cloned into a pSM vector containing the 3,527-nucleotide promoter of *egl-6* for HSN-specific degradation of ZF-tagged proteins^61^. All constructs were verified with sequencing.

### *C. elegans* CRISPR-Cas9 genome editing

Endogenous genome modifications were created by gonadal microinjection of CRISPR-Cas9 protein complexes^76^. Injection mixes consisted of 1.525 μM ALT-R Cas9, 5 μM tracer RNA and gRNA complexes (Integrated DNA Technologies), and either 5 μM single-stranded DNA (deletions) or 25–100 ng/μL PCR-amplified repair templates (insertions). ssDNA repair templates were designed with 75-base pair (bp) homology to each flank of the deletion. SYG-1(10YF) and SYG-1(7YF) repair templates were PCR-amplified from pNM286 and pNM434 vectors with ultramer oligonucleotides (Integrated DNA Technologies) containing 100bp of flanking genome homology. SYG-1-TurboID and SYG-1-FLAG-tev-SNAP construct repair templates were PCR-amplified from pNM207 and pNM366 vectors with ultramer oligonucleotides (Integrated DNA Technologies) containing 100bp of flanking genome homology. CRISPR–Cas9 repair templates either mutated the protospacer adjacent motif (PAM) site or altered four or more nucleotides within the gRNA binding site, in either case preserving amino-acid sequence. Guide RNAs were synthesized as ALT-R crRNAs (Integrated DNA Technologies). All genome edits were verified by PCR and sequencing.

### Microscopy and image processing

Images were obtained on a Nikon AX-R NSPARC super-resolution confocal system equipped with a 60X 1.46NA objective and 405, 488, 561, and 640nm lasers. HSN images were acquired from mid-L4 hermaphrodites, determined by vulval morphology. Quantification of synapse intensity was performed in Fiji/ImageJ on background-subtracted sum projections. Arbitrary unit intensity values were normalized to 100% for visualization in graphs. Representative HSN images are maximum intensity projections. FRAP assays were performed by bleaching a spot within each condensate with the corresponding excitation laser and monitoring recovery over time. FRAP curves were calculated after background subtraction and photobleaching correction and normalized to 100% for visualization.

### *In vivo* phosphosite identification

The SYG-1-FLAG-tev-SNAP strain was grown in 500 mL Complete S Basal liquid culture supplemented with OP50 *E. coli*. Worms were washed 3x with M9 buffer and snap frozen by dropwise addition to liquid nitrogen. Frozen droplets were ground to a fine powder with a cryogenic grinder (Cole-Parmer Freezer/Mill). The ground sample was thawed on ice and supplemented with equal volume of 2X RIPA buffer with protease and phosphatase inhibitors to yield a final 50 mM Tris pH 8, 150 mM NaCl, 1 mM EDTA, 1% SDS, 1% NP-40, 1 mM Na_3_VO_4_, 60 mM sodium β-glycerophosphate, 2.5 mM sodium pyrophosphate, 50 mM NaF, 1 mM PMSF, and 2 mM benzamidine. The sample was sonicated twice at 100% amplitude for 1 minute with cooling between steps (Fisherbrand 120 sonic dismembrator), followed by centrifugation at 4100 xg for 5 minutes to remove debris. Resultant lysates containing SNAP-tagged SYG-1 were incubated with SNAP-capture magnetic beads (NEB) overnight at 4°C to covalently bind. Beads were subsequently boiled for 10 minutes to denature all proteins, followed by extensive stringent washes to remove unbound material: two washes with 8 M urea-T buffer (8 M urea, 10 mM Tris pH 8.0, 0.2% Tween-20), two washes with strong detergent buffer (2% SDS, 1% sodium deoxycholate, 10 mM Tris pH 8.0), two washes with 1 M KCl-T (1 M KCl, 10 mM Tris pH 8.0), and five washes with TBS (10 mM Tris pH 8.0, 150 mM NaCl). During each wash, beads were incubated with rotation for 10 minutes before magnetic separation and beads were transferred to new tubes frequently.

A small sample was removed and cleaved with TEV protease at 16°C overnight to release the covalent linkage to the beads. This sample was run on a 4-12% Bis-tris gel (Invitrogen), transferred to a PVDF membrane, blotted with a primary mouse anti-FLAG antibody (Thermo Scientific FG4R) and a secondary goat anti-mouse antibody (Thermo Scientific 35518), and imaged on a Licor Odyssey CLx scanner.

Bead-bound samples were reduced, alkylated, and digested with AspN+GluC overnight before phosphopeptide enrichment. Both titanium oxide (TiO_2_) and iron (Fe-IMAC) enrichments were performed to recover phosphopeptides. First, 1mg Titansphere Phos-TiO Tip (GL Sciences) enrichment was performed according to the manufacturer’s protocol. Unbound flowthrough and washes were collected and desalted with GL-Tip SDB columns (GL Sciences) before further enrichment with homemade Fe-IMAC microcolumns as described^77^. Elutions from both enrichments were combined.

Peptide separations were performed using a Thermo Scientific Vanquish Neo UHPLC system operated in direct injection mode. Samples were loaded onto a 50 cm capillary column with an internal diameter of 75 µm. Separation was carried out at a constant flow rate of 0.300 µL/min using a 60-minute step gradient ranging from 1% acetonitrile to 65% acetonitrile.

Mass spectrometry data were acquired using an Orbitrap Exploris 480 mass spectrometer equipped with a nano-electrospray ionization (NSI) source operating in positive ion mode. Data acquisition was performed in Data-Dependent Acquisition (DDA) mode with a 3-second cycle time between master scans. Full MS1 scans were acquired in profile mode over a scan range of m/z 350–1500 at an Orbitrap resolution of 90,000. Precursors with charge states 2–6 and an intensity threshold exceeding 8,000 were selected for higher-energy collisional dissociation (HCD). Precursor ions were isolated using a 4 m/z isolation window and fragmented at a normalized collision energy (NCE) of 30%. MS2 scans were acquired in centroid mode in the Orbitrap at a resolution of 30,000 with a defined first mass of m/z 120. The MS2 normalized AGC target was 300% with automatic injection time. Dynamic exclusion was enabled to exclude precursors for 10 s following 1 selection event.

Raw LC-MS/MS files were processed in Thermo Scientific Proteome Discoverer (version 3.2.0.450) using a hybrid search algorithm strategy for phosphoproteomic analysis. Database searches were performed concurrently using Sequest HT and MS Amanda 3.0 against a custom FASTA database (Syg1.fasta) combined with common contaminants (PD_Contaminants_2015_5.fasta). Search parameters specified AspN+GluC cleavage with up to 3 missed cleavages, a precursor mass tolerance of 20 ppm, and a fragment mass tolerance of 0.02 Da. Peptide-spectrum matches (PSMs) were validated using fixed-value threshold scoring, and phosphorylation localization site probabilities were calculated using IMP-ptmRS with a 75% site probability threshold. High-confidence PSMs were subsequently integrated in the consensus workflow using strict parsimony rules for protein grouping, peptide/protein FDR filtering, and automated functional annotation via the Proteome Discoverer Protein Annotation Server. Data have been deposited to the ProteomeXchange Consortium via the PRIDE partner repository with the dataset identifier PXD083086.

### TurboID-mediated proximity labelling

TurboID proximity labeling was performed as described^78,79^. SYG-1-TurboID and control strains were grown in Complete S Basal liquid culture supplemented with an *E. coli* MG1655 food source. 1 mM biotin was added for a final 2 hours at 25°C. Worms were washed 3x with M9 buffer and snap frozen by dropwise addition to liquid nitrogen. Frozen droplets were ground to a fine powder with a cryogenic grinder (Cole-Parmer Freezer/Mill). The ground sample was thawed on ice and supplemented with an equal volume of 2X RIPA buffer with protease inhibitors to yield a final 50 mM Tris pH 7.5, 150 mM NaCl, 1 mM EDTA, 1% SDS, 1% NP-40, 1 mM PMSF, and 2 mM benzamidine. The sample was sonicated twice at 100% amplitude for 1 minute with cooling between steps (Fisherbrand 120 sonic dismembrator), followed by centrifugation at 4100 xg for 5 minutes to remove debris. Excess biotin was removed from the lysate with Zeba desalting columns (Thermo Scientific) on a chromatography system (Cytiva Akta Go). Lysates were further passed over a Histrap column to deplete endogenously biotinylated proteins tagged with His_6_ in the AX7884 background used. Streptavidin beads (Thermo Scientific Streptavidin T1) were added to each sample and incubated at 4°C overnight. Beads were washed twice with 2% SDS wash buffer, once with TBS-T buffer (50 mM Tris pH 7.5, 150 mM NaCl, 0.2% Tween-20), twice with 1 M KCl-T wash buffer (1 M KCl, 50 mM Tris– HCl pH 7.5, 1 mM EDTA, 0.2% Tween-20), twice with 0.1 M Na_2_CO_3_-T buffer (0.1 M Na_2_CO_3_, 0.2% Tween-20, pH 11.5), twice with 2 M urea-T buffer (2 M urea, 10 mM Tris pH 8.0, 0.2% Tween-20) and five times with TBS buffer (50 mM Tris pH 7.5, 150 mM NaCl). During each wash, beads were incubated with rotation for 10 minutes before magnetic separation and beads were transferred to new tubes frequently.

Peptides were separated on a Bruker nanoElute2 system operating in one-column mode with a PepSep column (10 cm, 75 µM, 1.9 µM particle size) maintained at 40°C. Chromatographic elution was performed at a constant flow rate of 0.50 µL/min over a total acquisition time of 62.33 min. The LC system was online-coupled to a timsTOF HT mass spectrometer (Bruker Daltonics) controlled via timsControl (v7.0.0.4). Data-independent acquisition (dia-PASEF) was operated in positive ion mode with full-scan MS spectra acquired over m/z 100-1700 at a rate of 1.00 Hz (rolling average: 10). The dia-PASEF isolation window scheme spanned a precursor mass range of m/z 460-1000 and an ion mobility range of −1/K0 ∼ 0.88 to 1.20 V.s/Cm^2^, organized across three diagonal m/z −1/K0 stepped bands (460∼660, 670∼870, and 880∼1000 m/z). Key TIMS tune parameters included: Funnel 1 RF at 300.0 Vpp, Funnel 2 RF at 200.0 Vpp, Multipole RF at 200.0 Vpp, Low Mass cutoff at 200.00 m/z, Collision RF at 1500.0 Vpp, Transfer Time of 60.0 µs, Pre-Pulse Storage time of 12.0µs, Ion Energy at 5.0 eV, and default Collision Energy at 10.0 eV. Spectra were saved in line and profile mode with mass calibration referenced against Tuning Mix ES-TOF (In-Batch).

Data-independent acquisition (DIA) files were processed using Spectronaut (version 21.0.260703.94842) operating in directDIA mode (directDIA+ Deep workflow). Database searches were performed using the Pulsar search engine against the uniprotkb_c_elegans_AND_model_organism_2025_07_31 protein database containing 29,021 protein entries. Digestion was specified for Trypsin/P, allowing up to 2 missed cleavages and peptide lengths ranging from 7 to 52 amino acids. The search strategy was configured to PTM Probing Search with a maximum allowance of 3 variable modifications per peptide. Identification cutoffs were set to a false discovery rate (FDR) Q-value of 0.01 at both the precursor level and experiment-wide protein level, alongside a run-level protein Q-value cutoff of 0.05. Precursor PEP and protein PEP cutoffs were configured to 0.2 and 0.75, respectively. Single hit protein identification followed a stratified single hit protein FDR rule defined by stripped sequence. Decoys were generated dynamically using a mutated strategy with neural network-predicted fragment sources. Protein inference was performed automatically using the IDPicker algorithm.

Quantification was calculated at the MS1 level using peak area integration and included ion mobility peak picking. Background noise removal was applied alongside interference correction (requiring a minimum of 2 MS1 and 3 MS2 ions). Peptides were grouped by stripped sequence, and protein groups were defined by Protein Group ID. Major group quantities were calculated as the mean peptide quantity using a Top N strategy (minimum 1, maximum 3). Cross-run normalization was not performed. Differential abundance testing between conditions was calculated using an unpaired t-test without assuming equal variance, using a Q-value threshold of 0.05 and a Log2 ratio candidate cutoff of 0.58. PTM localization was enabled with a minimum probability cutoff of 0.75. Data have been deposited to the ProteomeXchange Consortium via the PRIDE partner repository with the dataset identifier PXD083086.

GO analysis was performed with WormEnrichr^80,81^ using the list of significant (Q-value < 0.05 and Log2(TurboID:Control ratio) > 1) proteins.

### Expression and purification of recombinant proteins

Recombinant proteins were produced in Rosetta2(DE3) *E. coli* cells. Cultures were grown in TB medium to log phase (OD_600_ = 1.5) and induced with 0.4 mM isopropyl β-d-1-thiogalactopyranoside (IPTG) overnight at 18 °C. Cells were lysed in 50 mM Tris pH 8, 150 mM NaCl, 5 mM β-mercaptoethanol, and 0.1 mM phenylmethylsulfonyl fluoride (PMSF). Proteins to be labeled with NHS-ester fluorophores instead used 50 mM sodium phosphate pH 8 buffer for compatibility. Lysates were spun at 50,000 xg to pellet insoluble fractions. Proteins were first purified by affinity chromatography using Histrap or MBPtrap columns (Cytiva), cleaved by TEV protease at 16°C overnight if necessary, and polished with a Superdex Increase 200 or Superdex Increase 75 size-exclusion column (Cytiva). Proteins were concentrated with centrifugal protein concentrators with molecular weight cutoffs of 3,000 or 10,000 (Thermo Scientific). Proteins were labeled with NHS-ester dyes (Lumiprobe Alexa-488, Alexa-546, or Alexa-647 NHS-Ester) for 1h at room temperature, followed by dialysis to remove excess dye.

SYG-1 intracellular tail constructs were found to be relatively insoluble and required a modified procedure. After lysis and centrifugation performed as above, pellet fractions were recovered and re-solubilized in lysis buffer supplemented with 6M guanidine hydrochloride and 10% glycerol. Solubilized samples were bound to a HisTrap column and refolded on-column into 20 mM sodium phosphate pH 8, 150 mM NaCl, 5 mM β-mercaptoethanol, and 10% glycerol. Protein was eluted with 500 mM imidazole and subsequently polished with SEC as above, including 10% glycerol.

### Kinase assays

*In vitro* kinase assays consisted of 1 μg substrate, 1 μg SRC-1/SRC-2 kinase, 0.1 mM ATP, and 2 μCi γ^32^P-ATP (Revvity) in 40 mM Tris (pH 7.5), 20 mM magnesium chloride, 2 mM manganese chloride, 25 μM sodium orthovanadate, and 50 μM DTT. The samples were incubated at 30 °C for 45 min. Samples were quenched with LDS sample buffer and run on a 12% Bis-tris gel and stained with Coomassie SimplyBlue (Invitrogen) before drying. Dried gels were assembled in cassettes with a BAS-IP Phosphor screen (Cytiva) overnight and imaged on a Typhoon FLA 9500 system (GE Healthcare Life Sciences) with 635 nm excitation. Kinase assays upon supported lipid bilayers used 1 uM each SRC-1 and SRC-2 kinases in the buffer above, minus radiolabeled ATP, and were incubated for a minimum of 30 minutes at room temperature.

Mapping of phosphosites *in vitro* was performed by MS Bioworks. Phosphorylated SYG-1 tail was cut out of a gel, reduced with 10 mM dithiothreitol (DTT), alkylated with 50 mM iodoacetamide, and digested with trypsin at 37°C for 4 hours (Promega). Samples were analyzed by nano LC-MS/MS on a Waters M-Class HPLC coupled to a Thermo Fisher Orbitrap Fusion Lumos mass spectrometer. Peptides were loaded on a trapping column and eluted over a 75 μm analytical column at 350 nL/ min. Both columns were packed with Luna C18 resin (Phenomenex). The mass spectrometer was operated in data-dependent mode, with the Orbitrap operating at 60,000 FWHM and 15,000 FWHM for MS and MS/MS, respectively. APD was enabled and the instrument was run with a 3 s cycle for MS and MS/MS. The dataset is available at MassIVE (MSV000102872).

### In vitro binding assays

1 ug of purified SYG-1 tail was bound to 10 uL Ni-IMAC Resin (Thermo Scientific) in kinase buffer (25 mM Tris pH 8, 0.5 mM ATP, 20 mM magnesium chloride, 2 mM manganese chloride, and 50 μM DTT). In select samples, 1 ug of both SRC-1 and SRC-2 kinases were added. Samples were incubated overnight at 23°C to phosphorylate. 1 ug of each purified SH2 domain were subsequently added to bind for 2 hours. Three washes were performed to remove unbound material, and samples were run on 4-12% Bis-tris gels and stained with Simplyblue safestain (Thermo Scientific).

### Supported lipid bilayers

Supported lipid bilayers (SLBs) were formed on extensively cleaned 22 mm square glass coverslips (VWR)^82^. Coverslips were pre-cleaned with sequential sonication in a 2% Hellmanex III solution, ddH_2_O, 100% ethanol, and again in ddH_2_O. Further cleaning was performed by incubating for 10 minutes in a 1 M potassium hydroxide and 7.5% H_2_O_2_ solution at 70°C, rinsing with ddH2O, incubating for 10 minutes in a 2 M HCl and 7.5% H_2_O_2_ solution at 70°C, rinsing with ddH_2_O, and finally drying the coverslips in a nitrogen stream. A lipid mix composed of 97.5% DOPC, 2% DGS-NTA, 0.5% DOPE-PEG5000 (Avanti Research) was dried under vacuum and rehydrated with SLB buffer (25 mM Tris pH 7.5, 300 mM KCl, 1 mM MgCl_2_) at a final 1 mg/mL lipid concentration. Vigorous vortexing was performed to obtain multilamellar vesicles, which were subjected to five freeze-thaw cycles followed by extrusion 11x through 100 nm filters (Avanti Research) to obtain small unilamellar vesicles. Imaging chambers were created from 200 uL PCR tubes cut on top and bottom attached to the cleaned coverslips with UV glue (Thorlabs). SLBs were formed by the addition of 30 uL liposome mix and 1 uM CaCl_2_ to the chambers at 37°C for 30 minutes. Excess liposomes were removed and the buffer was adjusted with five 150 uL washes of 25 mM Tris pH 7.5, 150 mM NaCl, keeping the SLB hydrated under buffer the entire time. SYG-1 tail was added to SLBs at 100 nM for 30 minutes to bind. Excess was removed with five 150 uL washes. 2X concentrated samples of adapter protein combinations were prepared and added to the SYG-1 tail-bound SLBs to result in 1 uM final concentrations, followed by immediate imaging.

### Statistics and reproducibility

Standard one-way ANOVA tests with post-hoc Tukey multiple comparisons tests were performed on HSN intensity data in GraphPad Prism 9. Bar graphs depict all data points, means, and 95% confidence interval error bars. Each data point is a measurement from a separate animal. All imaging data were confirmed with a minimum of two imaging sessions. Two biological replicates were performed for *in vivo* phosphosite identification. TurboID was performed with three biological replicates for both experimental and control strains.

## Acknowledgements

We thank the de Bono lab for the gift of the AX7884 strain, members of the McDonald lab for critical reading of the manuscript, and Ludyanna Lebon and Dr. Rakesh Singh for their expertise and support with mass spectrometry experiments. Some strains were provided by the *Caenorhabditis* Genetics Center (CGC), which is funded by NIH Office of Research Infrastructure Programs P40 OD010440. Mass spectrometry experiments were performed in the Georgia Tech Systems Mass Spectrometry Core, supported by NIH S10 OD038327. This work was supported by NIH R00 NS123233 to N.A.M. The funders had no role in study design, data collection and analysis, decision to publish, or preparation of the manuscript.

## Author contributions

Conceptualization: Shenghan Wu and Nathan A. McDonald.

Funding acquisition: Nathan A. McDonald.

Investigation: Shenghan Wu, Nicole A. Morales, Daniel R. Li, and Nathan A. McDonald.

Methodology: Shenghan Wu and Nathan A. McDonald.

Project administration: Nathan A. McDonald.

Supervision: Nathan A. McDonald.

Writing – original draft: Shenghan Wu and Nathan A. McDonald.

Writing – review & editing: Shenghan Wu, Nicole A. Morales, Daniel Li, and Nathan A. McDonald.

---

**Figure S1.**
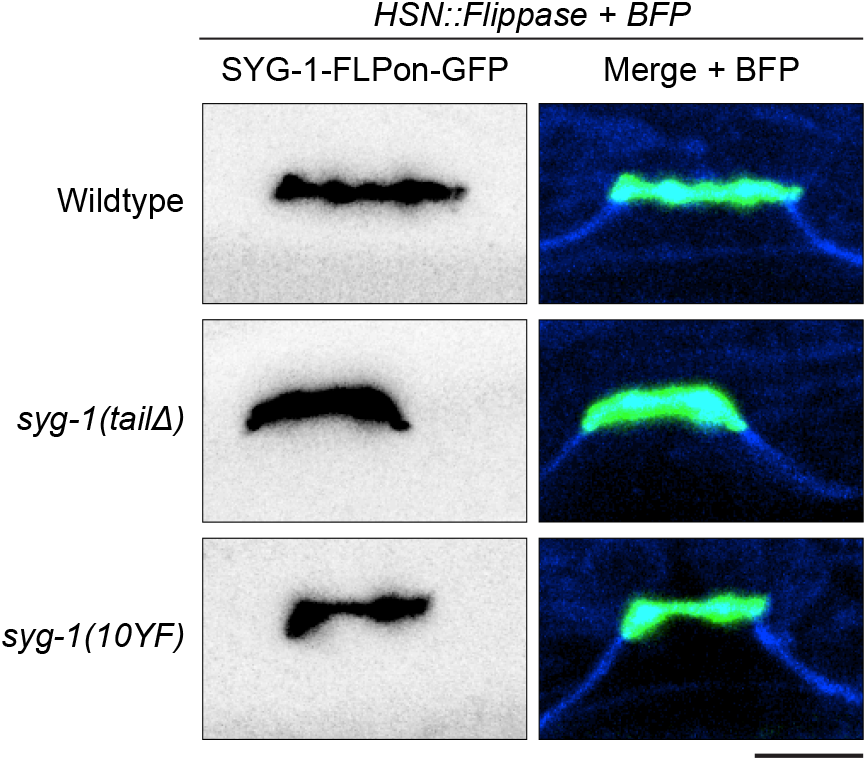
SYG-1 cytoplasmic tail mutants retain localization at the HSN synaptic zone. Confocal microscopy images of endogenous GFP-tagged SYG-1 mutants. BFP is expressed cytoplasmically. Scale bar, 5 μm.

**Figure S2.**
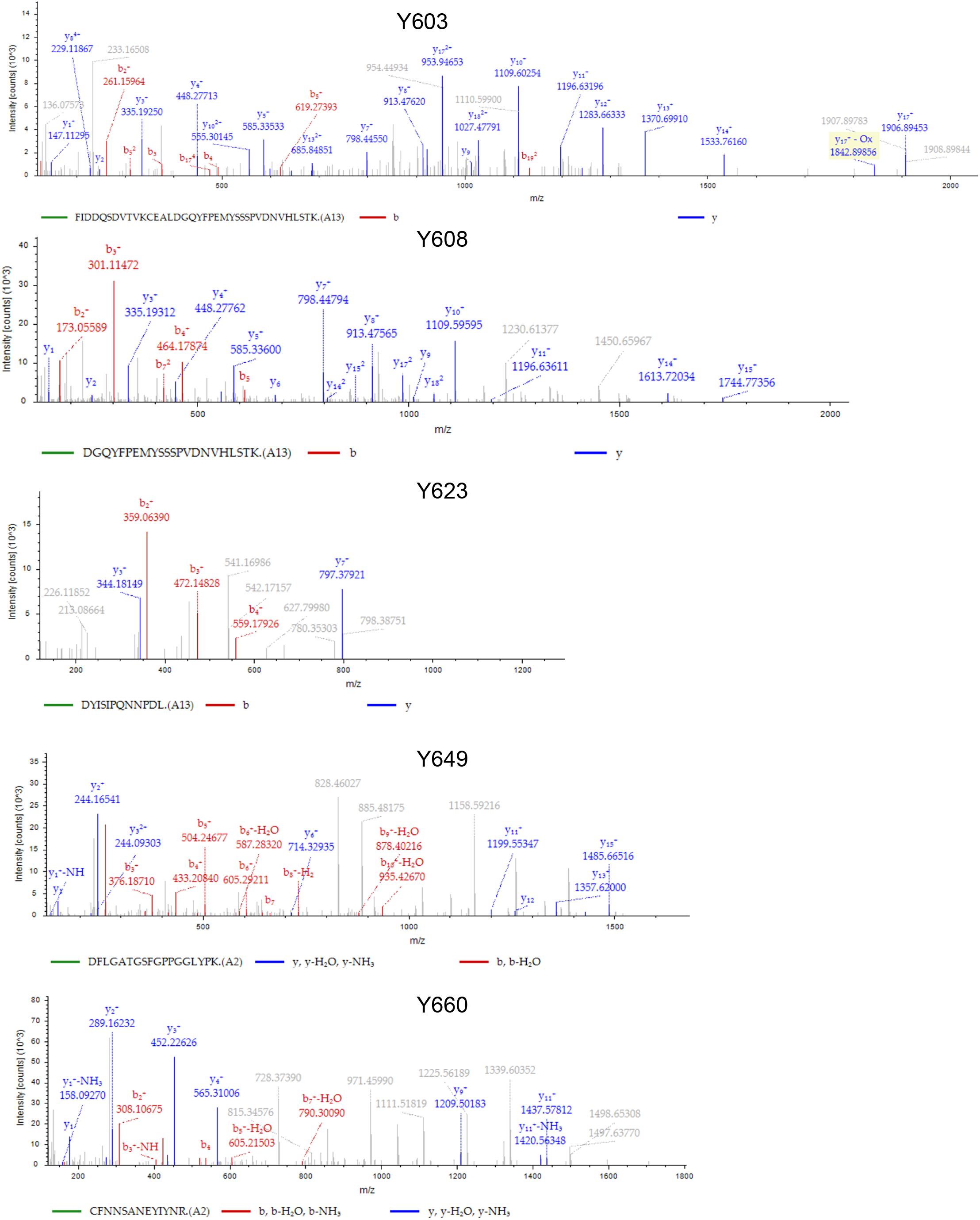
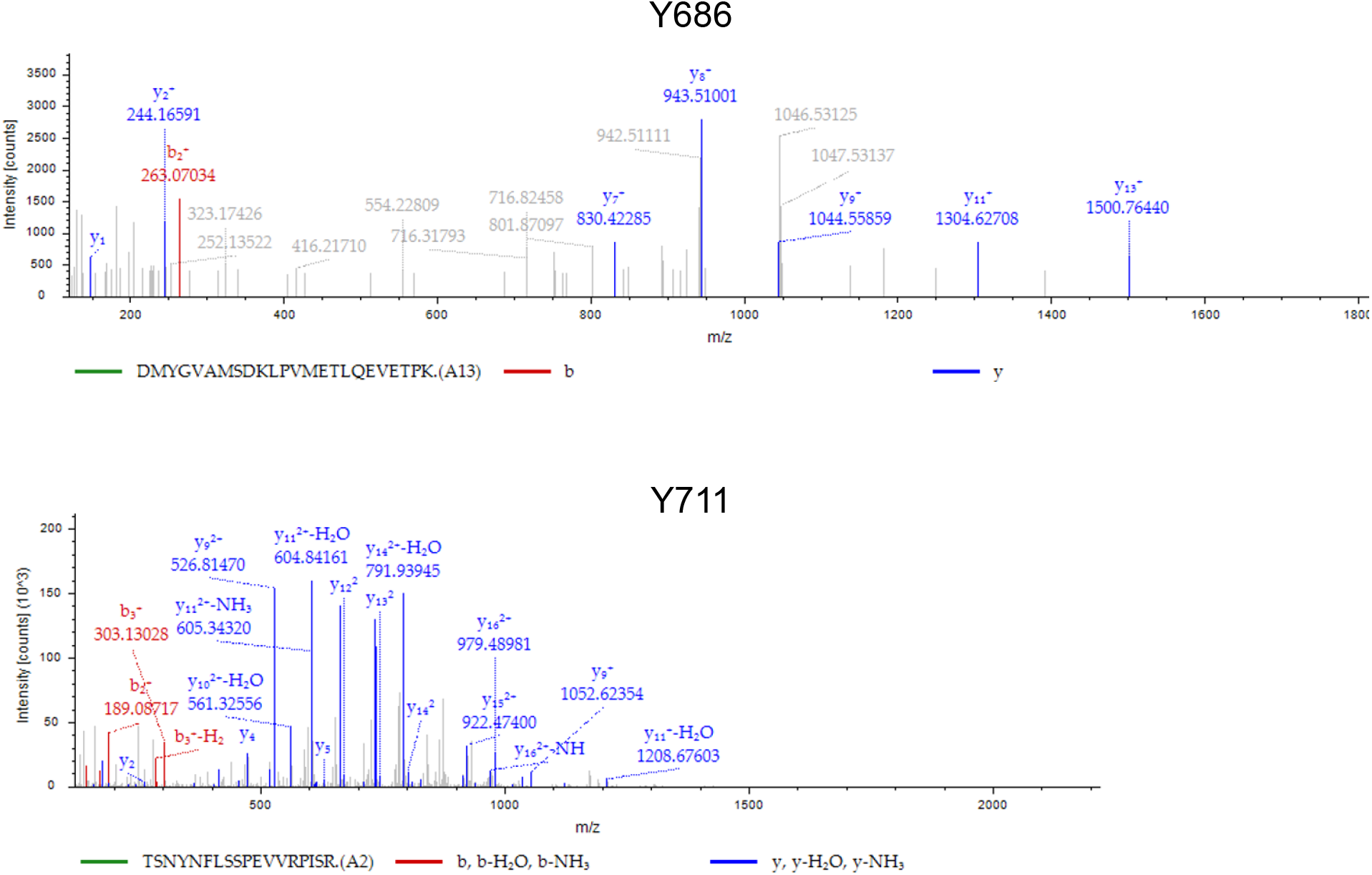
SYG-1 cytoplasmic tail *in vivo* phosphotyrosine spectra. Spectra of seven sites of tyrosine phosphorylation from a SYG-1-SNAP pulldown (Figure 2A-B and Table S1).

**Figure S3.**
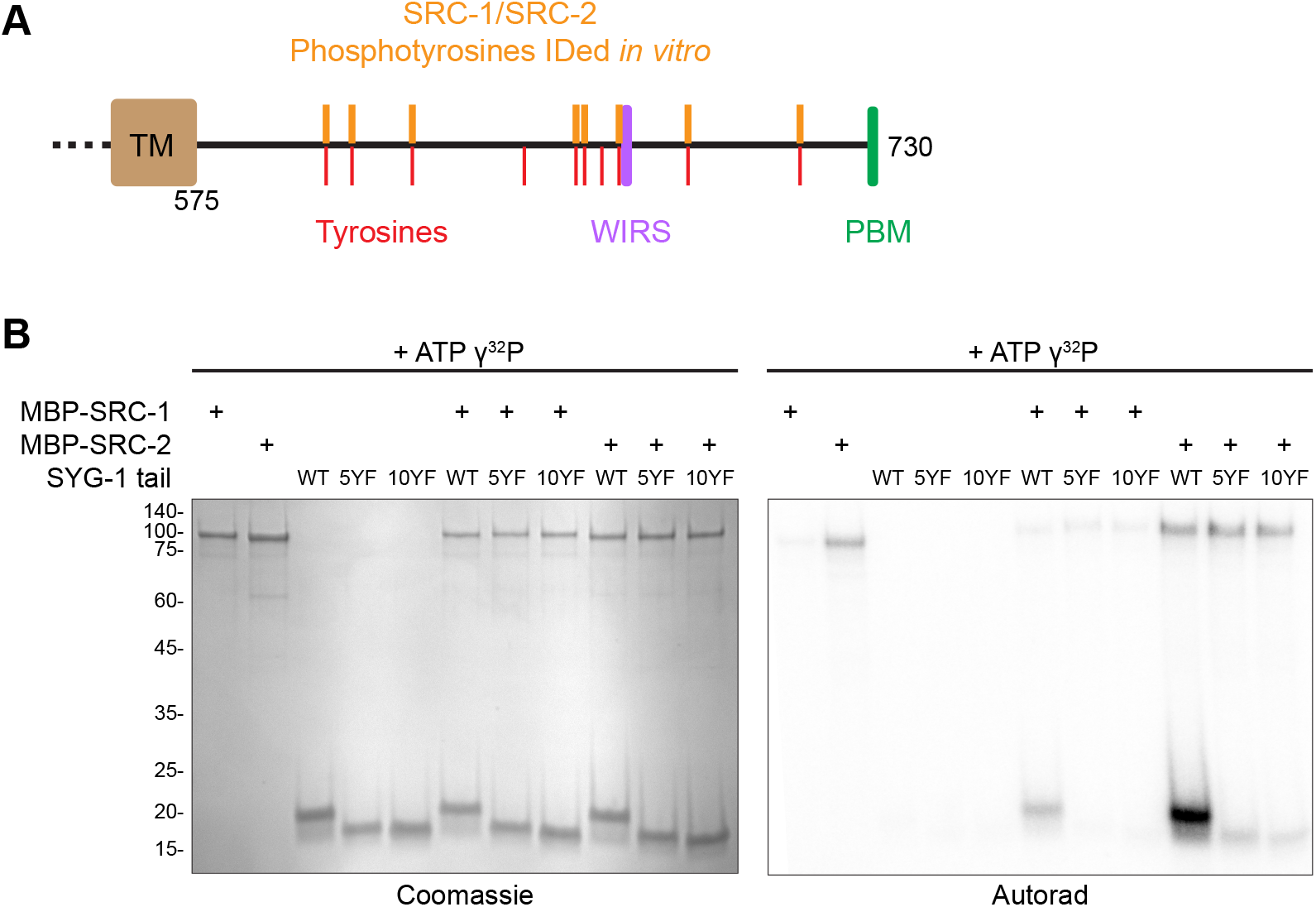
SRC-1 and SRC-2 tyrosine phosphorylate the SYG-1 cytoplasmic tail *in vitro*. A) Schematic of SYG-1 intracellular tail with phosphotyrosines identified in the *in vitro* kinase assay (Figure 4C) by mass spectrometry. See Table S3 for full data. B) *In vitro* kinase assay between purified recombinant SRC-1, SRC-2, and the indicated SYG-1 tail mutants.

**Figure S4.**
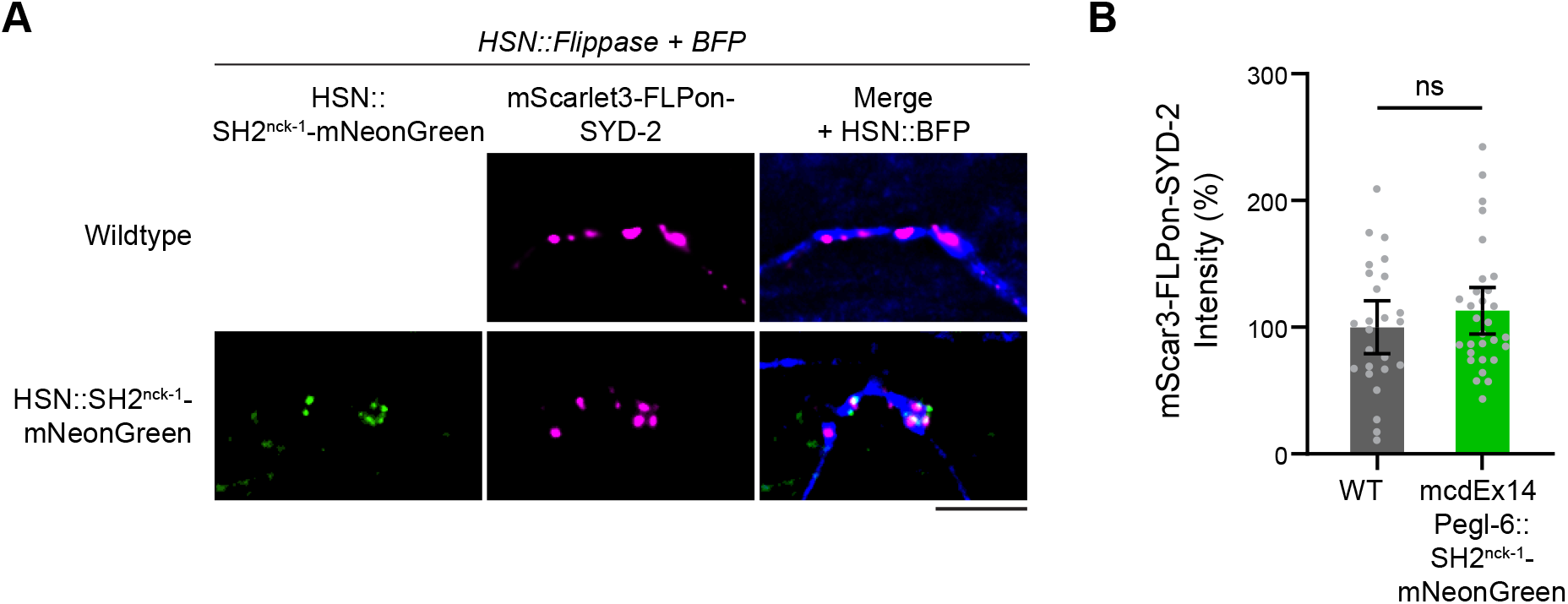
A SYG-1 phosphotyrosine sensor does not impact synapse formation. A) HSN synapse formation phenotypes with or without SH2^nck-1^-mNeonGreen expression, visualized with an endogenously-tagged SYD-2 active zone marker. Scale bar, 5 μm. B) Quantification of HSN synaptic marker fluorescent intensities in (B). ****, p < 0.0001.

**Figure S5.**
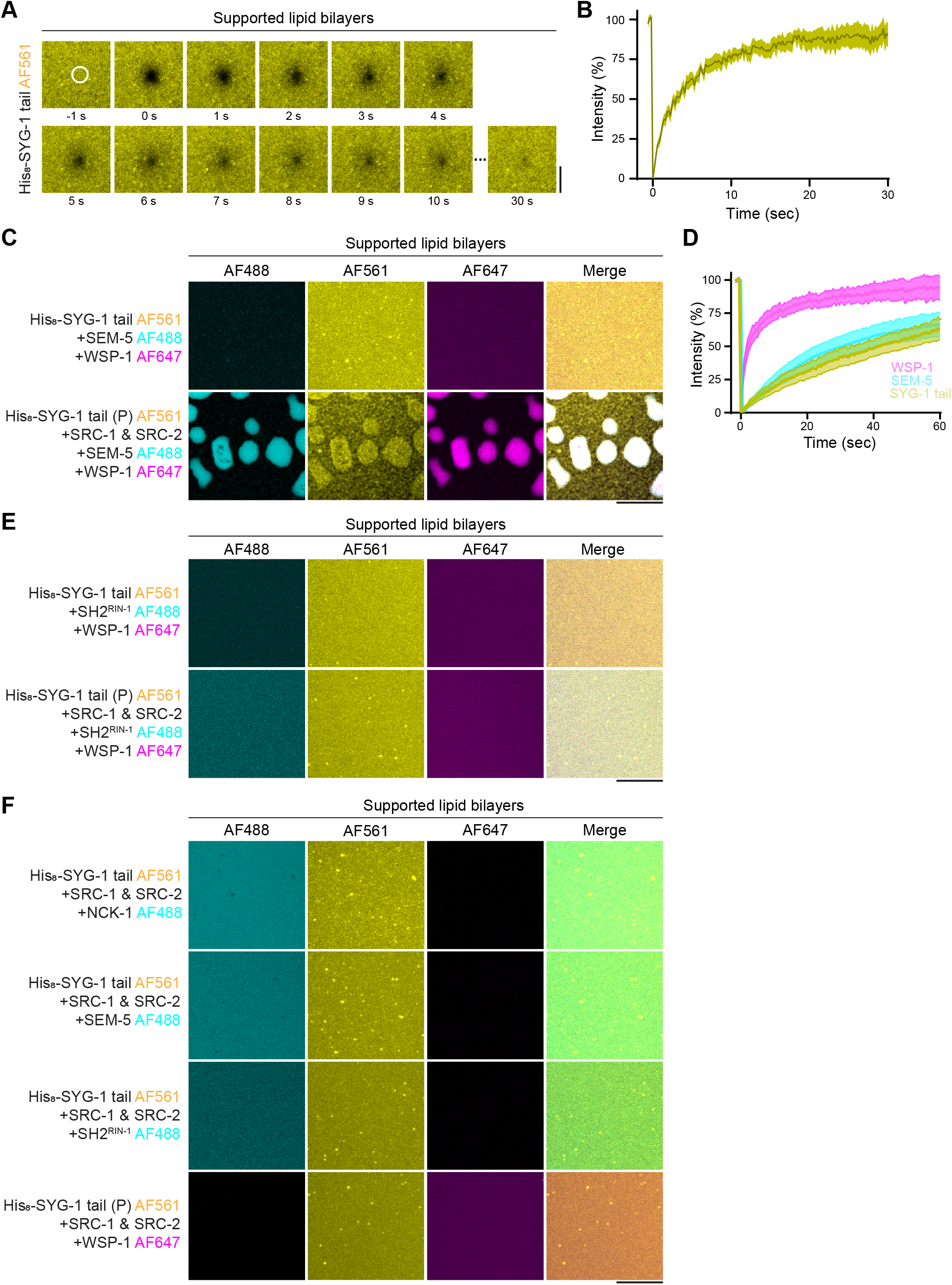
Additional supported lipid bilayer conditions. A-B) Fluorescence recovery after photobleaching (FRAP) of supported lipid bilayer-bound SYG-1 tail. The indicated white circular region was bleached and recovery was monitored. Robust recovery was observed, indicating fluid bilayers have been produced. C) Addition of SEM-5 and WSP-1 to the SYG-1 tail-bound SLBs in the presence or absence of SRC-1/2 phosphorylation. D) Fluorescence recovery after photobleaching (FRAP) of each condensate component from (C) reveals liquid dynamics within condensates. E) Addition of SH2^RIN-1^ and WSP-1 to the SYG-1 tail-bound SLBs with or without SRC-1/2 phosphorylation. F) Individual addition of NCK-1, SEM-5, SH2^RIN-1^, or WSP-1 to the SYG-1 tail-bound SLBs with SRC-1/2 phosphorylation. No condensates were observed. Scale bars, 10 um.

**Table S1. SYG-1 in vivo phosphopeptides.**

**Table S2. SYG-1-TurboID proximity labeling.**

**Table S3. Sites of SRC-1 and SRC-2 phosphorylation on the SYG-1 tail *in vitro*.**

**Table S4. *C. elegans* strains used in this study.**

**Table S5. Plasmids used in this study.**

## References

1. Südhof, T. C. The cell biology of synapse formation. J. Cell Biol. 220, 1–18 (2021).

2. Emperador-Melero, J. & Kaeser, P. S. Assembly of the presynaptic active zone. Curr. Opin. Neurobiol. 63, 95–103 (2020).

3. Südhof, T. C. The Presynaptic Active Zone. Neuron 75, 11–25 (2012).

4. Sheng, M. & Kim, E. The Postsynaptic Organization of Synapses. Cold Spring Harb. Perspect. Biol. 3, a005678 (2011).

5. Chen, X., Wu, X., Wu, H. & Zhang, M. Phase separation at the synapse. Nat. Neurosci. 23, 301–310 (2020).

6. Banani, S. F., Lee, H. O., Hyman, A. A. & Rosen, M. K. Biomolecular condensates: organizers of cellular biochemistry. Nat. Rev. Mol. Cell Biol. 18, 285–298 (2017).

7. Alberti, S. & Hyman, A. A. Biomolecular condensates at the nexus of cellular stress, protein aggregation disease and ageing. Nat. Rev. Mol. Cell Biol. 22, 196–213 (2021).

8. McDonald, N. A., Fetter, R. D. & Shen, K. Assembly of synaptic active zones requires phase separation of scaffold molecules. Nature 588, 454–458 (2020).

9. Emperador-Melero, J. et al. PKC-phosphorylation of Liprin-α3 triggers phase separation and controls presynaptic active zone structure. Nat. Commun. 12, 3057 (2021).

10. Jin, G. et al. The liprin-α/RIM complex regulates the dynamic assembly of presynaptic active zones via liquid–liquid phase separation. PLOS Biol. 23, e3002817 (2025).

11. Wu, X. et al. RIM and RIM-BP Form Presynaptic Active-Zone-like Condensates via Phase Separation. Mol. Cell 73, 971–984.e5 (2019).

12. Milovanovic, D., Wu, Y., Bian, X. & De Camilli, P. A liquid phase of synapsin and lipid vesicles. Science 361, 604–607 (2018).

13. Park, D. et al. Cooperative function of synaptophysin and synapsin in the generation of synaptic vesicle-like clusters in non-neuronal cells. Nat. Commun. 12, 263 (2021).

14. Feng, Z., Chen, X., Zeng, M. & Zhang, M. Phase separation as a mechanism for assembling dynamic postsynaptic density signalling complexes. Curr. Opin. Neurobiol. 57, 1–8 (2019).

15. Chen, S. et al. Native postsynaptic density is a functional condensate formed via phase separation. Cell Rep. 45, 116723 (2026).

16. Lv, P. et al. O-GlcNAcylation modulates liquid–liquid phase separation of SynGAP/PSD-95. Nat. Chem. 1–10 (2022) doi:10.1038/s41557-022-00946-9.

17. Jia, B. et al. Shank3 oligomerization governs material properties of the postsynaptic density condensate and synaptic plasticity. Cell 0, (2025).

18. Verpoort, B. & Wit, J. de. Cell Adhesion Molecule Signaling at the Synapse: Beyond the Scaffold. Cold Spring Harb. Perspect. Biol. 16, a041501 (2024).

19. Gomez, A. M., Traunmüller, L. & Scheiffele, P. Neurexins: molecular codes for shaping neuronal synapses. Nat. Rev. Neurosci. 22, 137–151 (2021).

20. Bemben, M. A., Shipman, S. L., Nicoll, R. A. & Roche, K. W. The cellular and molecular landscape of neuroligins. Trends Neurosci. 38, 496–505 (2015).

21. Young, T. R. & Leamey, C. A. Teneurins: Important regulators of neural circuitry. Int. J. Biochem. Cell Biol. 41, 990–993 (2009).

22. Südhof, T. C. Signaling by latrophilin adhesion-GPCRs in synapse assembly. Neuroscience 10.1016/j.neuroscience.2025.03.041 (2025) doi:10.1016/j.neuroscience.2025.03.041.

23. Takahashi, H. & Craig, A. M. Protein tyrosine phosphatases PTPδ, PTPσ, and LAR: presynaptic hubs for synapse organization. Trends Neurosci. 36, 522–534 (2013).

24. Hruska, M. & Dalva, M. B. Ephrin regulation of synapse formation, function and plasticity. Mol. Cell. Neurosci. 50, 35–44 (2012).

25. Sytnyk, V., Leshchyns’ka, I. & Schachner, M. Neural Cell Adhesion Molecules of the Immunoglobulin Superfamily Regulate Synapse Formation, Maintenance, and Function. Trends Neurosci. 40, 295–308 (2017).

26. Zinn, K. & Özkan, E. Neural immunoglobulin superfamily interaction networks. Curr. Opin. Neurobiol. 45, 99–105 (2017).

27. Takahashi, H. et al. Postsynaptic TrkC and Presynaptic PTPσ Function as a Bidirectional Excitatory Synaptic Organizing Complex. Neuron 69, 287–303 (2011).

28. Choi, Y. et al. SALM5 trans-synaptically interacts with LAR-RPTPs in a splicing-dependent manner to regulate synapse development. Sci. Rep. 6, 26676 (2016).

29. Takahashi, H. et al. Selective control of inhibitory synapse development by Slitrk3-PTPδ trans-synaptic interaction. Nat. Neurosci. 15, 389–398 (2012).

30. Woo, J. et al. Trans-synaptic adhesion between NGL-3 and LAR regulates the formation of excitatory synapses. Nat. Neurosci. 12, 428–437 (2009).

31. Cosmanescu, F. et al. Neuron-Subtype-Specific Expression, Interaction Affinities, and Specificity Determinants of DIP/Dpr Cell Recognition Proteins. Neuron 100, 1385–1400.e6 (2018).

32. Um, J. W. & Ko, J. LAR-RPTPs: synaptic adhesion molecules that shape synapse development. Trends Cell Biol. 23, 465–475 (2013).

33. Zhang, X., Chen, X., Matúš, D. & Südhof, T. C. Reconstitution of synaptic junctions orchestrated by teneurin-latrophilin complexes. Science 387, 322–329 (2025).

34. Wang, S. et al. Alternative splicing of latrophilin-3 controls synapse formation. Nature 626, 128–135 (2024).

35. Sando, R. & Südhof, T. C. Latrophilin GPCR signaling mediates synapse formation. eLife 10, e65717 (2021).

36. Collins, K. M. et al. Activity of the C. elegans egg-laying behavior circuit is controlled by competing activation and feedback inhibition. eLife 5, e21126 (2016).

37. Huang, Y.-C. et al. A single neuron in *C. elegans* orchestrates multiple motor outputs through parallel modes of transmission. Curr. Biol. 33, 4430–4445.e6 (2023).

38. White, J. G., Southgate, E., Thomson, J. N. & Brenner, S. The structure of the nervous system of the nematode Caenorhabditis elegans. Philos. Trans. R. Soc. Lond. B Biol. Sci. 314, 1–340 (1986).

39. Shen, K. & Bargmann, C. I. The Immunoglobulin Superfamily Protein SYG-1 Determines the Location of Specific Synapses in C. elegans. Cell 112, 619–630 (2003).

40. Shen, K., Fetter, R. D. & Bargmann, C. I. Synaptic Specificity Is Generated by the Synaptic Guidepost Protein SYG-2 and Its Receptor, SYG-. Cell 116, 869–881 (2004).

41. Özkan, E. et al. Extracellular Architecture of the SYG-1/SYG-2 Adhesion Complex Instructs Synaptogenesis. Cell 156, 482–494 (2014).

42. Chia, P. H., Chen, B., Li, P., Rosen, M. K. & Shen, K. Local F-actin Network Links Synapse Formation and Axon Branching. Cell 156, 208–220 (2014).

43. Hunter, T. The Genesis of Tyrosine Phosphorylation. Cold Spring Harb. Perspect. Biol. 6, a020644 (2014).

44. Chao, D. L. & Shen, K. Functional dissection of SYG-1 and SYG-2, cell adhesion molecules required for selective synaptogenesis in C. elegans. Mol. Cell. Neurosci. 39, 248–257 (2008).

45. McDonald, N. A., Tao, L., Dong, M.-Q. & Shen, K. SAD-1 kinase controls presynaptic phase separation by relieving SYD-2/Liprin-α autoinhibition. PLOS Biol. 21, e3002421 (2023).

46. Branon, T. C. et al. Efficient proximity labeling in living cells and organisms with TurboID. Nat. Biotechnol. 36, 880–887 (2018).

47. Roskoski, R. Src kinase regulation by phosphorylation and dephosphorylation. Biochem. Biophys. Res. Commun. 331, 1–14 (2005).

48. Roskoski, R. Src protein–tyrosine kinase structure and regulation. Biochem. Biophys. Res. Commun. 324, 1155–1164 (2004).

49. Pawson, T. & Gish, G. D. SH2 and SH3 domains: From structure to function. Cell 71, 359–362 (1992).

50. Pawson, T., Gish, G. D. & Nash, P. SH2 domains, interaction modules and cellular wiring. Trends Cell Biol. 11, 504–511 (2001).

51. Verma, R. et al. Nephrin ectodomain engagement results in Src kinase activation, nephrin phosphorylation, Nck recruitment, and actin polymerization. J. Clin. Invest. 116, 1346–1359 (2006).

52. Jones, N. et al. Nck adaptor proteins link nephrin to the actin cytoskeleton of kidney podocytes. Nature 440, 818–823 (2006).

53. Mayer, B. J. & Baltimore, D. Signalling through SH2 and SH3 domains. Trends Cell Biol. 3, 8–13 (1993).

54. Schlessinger, J. & Lemmon, M. A. SH2 and PTB Domains in Tyrosine Kinase Signaling. Sci. STKE 2003, re12–re12 (2003).

55. Case, L. B., Zhang, X., Ditlev, J. A. & Rosen, M. K. Stoichiometry controls activity of phase-separated clusters of actin signaling proteins. Science 363, 1093–1097 (2019).

56. Banjade, S. & Rosen, M. K. Phase transitions of multivalent proteins can promote clustering of membrane receptors. eLife 3, e04123 (2014).

57. Su, X. et al. Phase separation of signaling molecules promotes T cell receptor signal transduction. Science 352, 595–599 (2016).

58. Ditlev, J. A. et al. A composition-dependent molecular clutch between T cell signaling condensates and actin. eLife 8, e42695 (2019).

59. Huang, W. Y. C. et al. A molecular assembly phase transition and kinetic proofreading modulate Ras activation by SOS. Science 363, 1098–1103 (2019).

60. Jaqaman, K. & Ditlev, J. A. Biomolecular condensates in membrane receptor signaling. Curr. Opin. Cell Biol. 69, 48–54 (2021).

61. Armenti, S. T., Lohmer, L. L., Sherwood, D. R. & Nance, J. Repurposing an endogenous degradation system for rapid and targeted depletion of C. elegans proteins. Development 141, 4640–4647 (2014).

62. Wong, L. E. et al. Tripartite phase separation of two signal effectors with vesicles priming B cell responsiveness. Nat. Commun. 11, 848 (2020).

63. Garg, P., Verma, R., Nihalani, D., Johnstone, D. B. & Holzman, L. B. Neph1 Cooperates with Nephrin To Transduce a Signal That Induces Actin Polymerization. Mol. Cell. Biol. 27, 8698–8712 (2007).

64. Kim, S., Kalappurakkal, J. M., Mayor, S., Rosen, M. K. & Gardel, M. Phosphorylation of nephrin induces phase separated domains that move through actomyosin contraction. Mol. Biol. Cell 30, 2996–3012 (2019).

65. Sun, S., GrandPre, T., Limmer, D. T. & Groves, J. T. Kinetic frustration by limited bond availability controls the LAT protein condensation phase transition on membranes. Sci. Adv. 8, eabo5295 (2022).

66. Wu, X. et al. Vesicle Tethering on the Surface of Phase-Separated Active Zone Condensates. Mol. Cell 81, 13–24.e7 (2021).

67. Bachir, A. I., Horwitz, A. R., Nelson, W. J. & Bianchini, J. M. Actin-Based Adhesion Modules Mediate Cell Interactions with the Extracellular Matrix and Neighboring Cells. Cold Spring Harb. Perspect. Biol. 9, a023234 (2017).

68. Vicente-Manzanares, M. & Horwitz, A. R. Adhesion dynamics at a glance. J. Cell Sci. 124, 3923–3927 (2011).

69. Bingham, D. et al. Presynapses contain distinct actin nanostructures. J. Cell Biol. 222, e202208110 (2023).

70. Butler-Hallissey, C. & Leterrier, C. Super-resolution imaging of the neuronal cytoskeleton. Npj Imaging 2, 50 (2024).

71. Tumminia, S. et al. Deciphering the Nanoscale Architecture of Presynaptic Actin Using a Micropatterned Presynapse-on-Glass Model. J. Neurosci. 46, (2026).

72. Lei, W., Omotade, O. F., Myers, K. R. & Zheng, J. Q. Actin cytoskeleton in dendritic spine development and plasticity. Curr. Opin. Neurobiol. 39, 86–92 (2016).

73. Brodin, L. & Shupliakov, O. Actin cage for the synaptic vesicle liquid phase. Cell Rep. 44, (2025).

74. Nelson, J. C., Stavoe, A. K. H. & Colón-Ramos, D. A. The actin cytoskeleton in presynaptic assembly. Cell Adhes. Migr. 7, 379–387 (2013).

75. Chia, P. H., Patel, M. R. & Shen, K. NAB-1 instructs synapse assembly by linking adhesion molecules and F-actin to active zone proteins. Nat. Neurosci. 15, 234–242 (2012).

76. Ghanta, K. S. & Mello, C. C. Melting dsDNA Donor Molecules Greatly Improves Precision Genome Editing in Caenorhabditis elegans. Genetics 216, 643–650 (2020).

77. Sanford, E. J. & Smolka, M. B. Fe-NTA Microcolumn Purification of Phosphopeptides from Immunoprecipitation (IP) Eluates for Mass Spectrometry Analysis. Bio-Protoc. 11, (2021).

78. Artan, M., Hartl, M., Chen, W. & de Bono, M. Depletion of endogenously biotinylated carboxylases enhances the sensitivity of TurboID-mediated proximity labeling in Caenorhabditis elegans. J. Biol. Chem. 298, 102343 (2022).

79. Artan, M. et al. Interactome analysis of *Caenorhabditis elegans* synapses by TurboID-based proximity labeling. J. Biol. Chem. 297, 101094 (2021).

80. Chen, E. Y. et al. Enrichr: interactive and collaborative HTML5 gene list enrichment analysis tool. BMC Bioinformatics 14, 128 (2013).

81. Kuleshov, M. V. et al. Enrichr: a comprehensive gene set enrichment analysis web server 2016 update. Nucleic Acids Res. 44, W90–W97 (2016).

82. Cheng, X., Ullo, M. F. & Case, L. B. Reconstitution of Phase-Separated Signaling Clusters and Actin Polymerization on Supported Lipid Bilayers. Front. Cell Dev. Biol. 10, (2022).

